# Epigenetic control of metabolic identity across cell types

**DOI:** 10.1101/2024.07.24.604914

**Authors:** Maria Pires Pacheco, Déborah Gerard, Riley J. Mangan, Alec R. Chapman, Dennis Hecker, Manolis Kellis, Marcel H. Schulz, Lasse Sinkkonen, Thomas Sauter

## Abstract

**Background:** Constraint-based network modeling is a powerful genomic-scale approach for analyzing cellular metabolism, capturing metabolic variations across tissues and cell types, and defining the metabolic identity essential for identifying disease-associated transcriptional states.

**Results:** Using RNA-seq and epigenomic data from the EpiATLAS resource of the International Human Epigenome Consortium (IHEC), we reconstructed metabolic networks for 1,555 samples spanning 58 tissues and cell types. Analysis of these networks revealed the distribution of metabolic functionalities across human cell types and provides a compendium of human metabolic activity. This integrative approach allowed us to define, across tissues and cell types, i) reactions that fulfil the basic metabolic processes (core metabolism), and ii) cell type-specific functions (unique metabolism), that shape the metabolic identity of a cell or a tissue. Integration with EpiATLAS-derived cell-type-specific gene-level chromatin states and enhancer-gene interactions identified enhancers, transcription factors, and key nodes controlling core and unique metabolism. Transport and first reactions of pathways were enriched for high expression, active chromatin state, and Polycomb-mediated repression in cell types where pathways are inactive, suggesting that key nodes are targets of repression.

**Discussion:** This integrative analysis forms the basis for identifying regulation points that control metabolic identity in human cells.

## Background

Identifying the molecular hallmarks of cell identity is crucial for understanding cellular differentiation in development and disease and can inform strategies for cellular reprogramming. Although the concept of cell identity remains debated, it is often inferred from cellular functions and phenotypes governed by gene expression patterns maintained by identity-specifying transcription factors (TFs) ^1^.

Metabolism is a key determinant of cell identity. For example, in type II diabetes, high blood glucose levels can disrupt beta cell identity, resulting in loss of beta cell-specific gene expression and gain of gene expression programs of other islet cell types ^2^. Similarly, oxidative phosphorylation and reactive oxygen species (ROS) signalling drive the differentiation of intestinal stem cells into Paneth cells ^3^. The centrality of metabolism in cellular identity is further supported by the observation that culture media, rather than tissue of origin, governs pluripotent stem cell identity.

Metabolic identity extends the cellular identity concept to the activities of biochemical pathways in a cell. Metabolic identity includes the core metabolism, a set of biochemical reactions shared across cell types that maintain energy and chemical homeostasis, and unique metabolism, or cell type-specific metabolic processes in differentiated cells, such as alcohol detoxification in the liver and melanin production in skin melanocytes.

Metabolic network modelling, which integrates cell type-specific epigenomic maps with the biochemical reactome, is a powerful approach to capture metabolic identity across cell types and identity disruptions in disease states^4–10^. Metabolic models provide key insights into how the inactivation of rate-limiting enzymes controls metabolic flux, the interactions between metabolic pathways and compartments, and the ultimate capacity of cells to perform metabolic functions ^11^.

Cell-type-specific metabolism is achieved through precise regulation of pathway enzymes through the epigenetic control of gene expression, post-translational modifications, and compartmental sequestration of metabolites and co-factors. In support of this, we previously demonstrated that cells place high regulatory loads on rate-limiting pathway enzymes ^12^. Specifically, loci encoding transporter proteins that control substrate availability exhibit high H3K27ac signal in human macrophages ^13^.

More generally, cell-type-specific gene expression is maintained by regulatory elements such as promoters, enhancers, and silencers, whose activities have been mapped genome-wide through comprehensive ChIP-seq, ATAC-seq, and DNase-seq profiling of histone modifications and open chromatin generated by Roadmap Epigenomics ^14^, ENCODE ^15^, and other consortia. The recent International Human Epigenome Consortium EpiATLAS resource (International Human Epigenome Consortium, EpiATLAS - a reference for human epigenomic research, in preparation) ^16^ greatly expands these resources, providing an unprecedented opportunity to dissect the regulatory control of metabolism. Notably, enhancer-gene interactions inferred in the EpiATLAS resource using the generalized Activity-By-Contact (ABC) model ^17^ provide opportunities to connect regulatory elements, and their upstream transcription factor regulators, to metabolic genes.

Probabilistic graphical model methods, including ChromHMM ^18^ and Segway ^19^, infer cell type-specific chromatin states from combinatorial and spatial patterns of epigenomic signals. While ChromHMM and Segway define chromatin states at the level of individual loci, the recent ChromGene ^20^ method defines gene-level chromatin state by integrating histone modification signals throughout the gene body and flanking regions using a mixture of hidden Markov models ^21^. ChromGene defines six active states (1-6), one poised state (7), and five repressive states (8-12) **(Supplementary Table 1)**, based on combinations of ten histone modifications. This gene-level view provides better insight into which genes encoding for enzymes and transporters in the metabolic networks are active at the chromatin level, and which are transcriptionally repressed.

In this work, we reconstructed metabolic models across 1,555 biosamples from the IHEC EpiATLAS resource spanning 58 human cell types and tissues to identify core and tissue-specific metabolic reactions. We leveraged EpiATLAS-derived enhancer-gene interactions to discover regulatory elements and upstream transcription factors controlling these processes. We observe that transporter genes are enriched for cell type-specific regulatory control and that many core metabolic pathways expressed across tissue types exhibit cell-type-specific regulatory control, providing a mechanistic basis for tissue-specific metabolic reprogramming of core pathways.

## Results

### Metabolic models capture core and tissue-specific metabolism

We used RNA-seq data of 1,555 samples from the EpiATLAS (International Human Epigenome Consortium, EpiATLAS - a reference for human epigenomic research, in preparation) ^16^ resource to reconstruct metabolic models that served as scaffolds to map gene-level chromatin states

(ChromGene) **(Supplementary Table 1)** and enhancer activity data from the corresponding samples to the human metabolic network to define: 1) the core metabolic biochemical reactions across cell types and tissues, 2) the metabolic identity for individual cell types and tissues, and 3) the regulatory mechanisms controlling this identity **(Figure 1)**.

**Figure 1:**
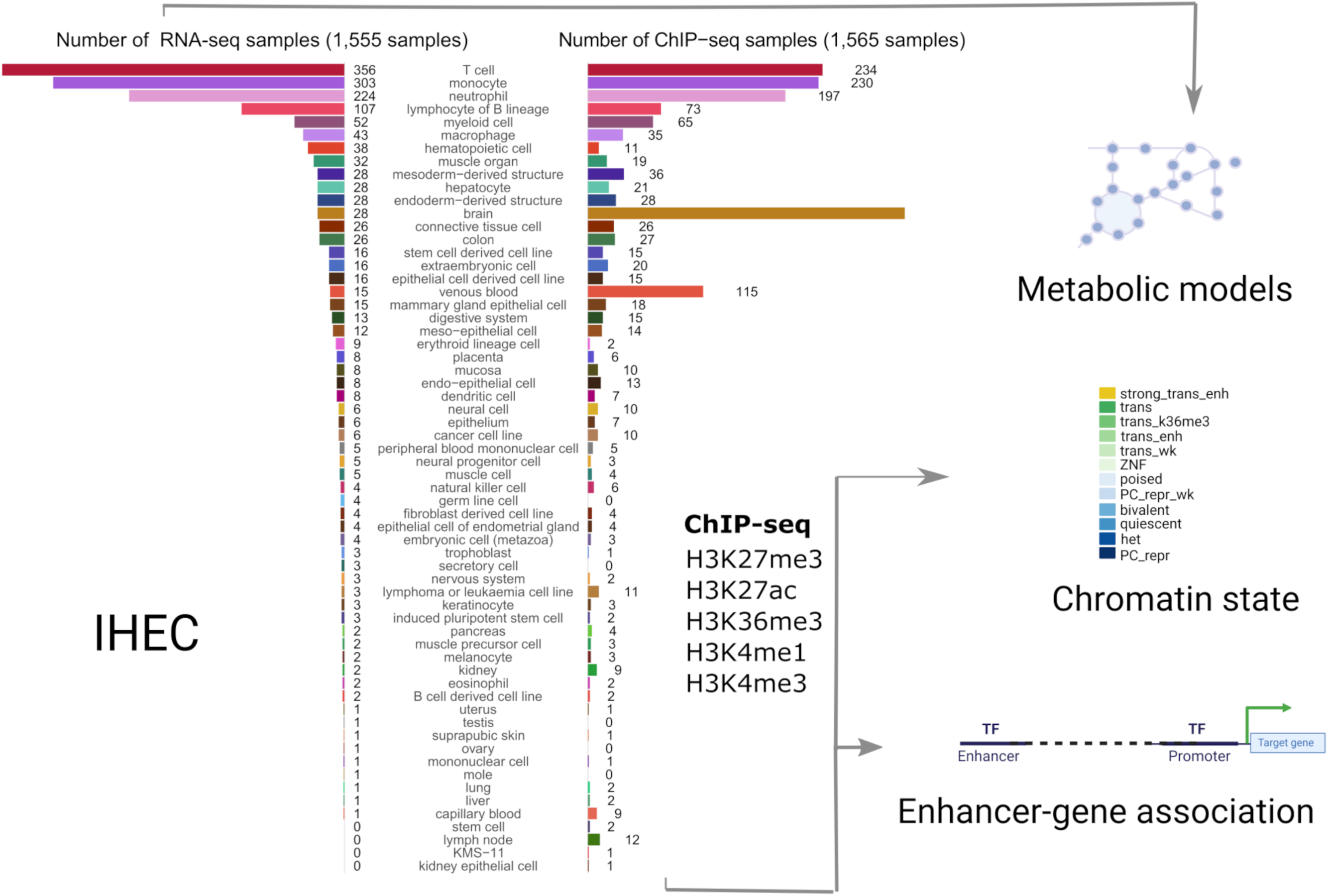
**Systematic modelling of epigenetic control of metabolism**. The number of integrated RNA-seq and ChIP-seq samples for each tissue and cell type is represented as bar plots. We used RNA-seq profiles (left bar plot) from 58 tissues and cell types to reconstruct metabolic networks with rFASTCORMICS ^5^. These models were then used as scaffolds to map enhancer-activity scores (1,565 samples across 56 cell types and tissues, right bar plot) as well as gene-level ChromGene chromatin state annotations (1,698 samples across 57 cell types and tissues, not shown as samples distribution is similar to the enhancer activity scores) to identify key regulation points. 12 ChromGene states were defined based on the integration of the chromatin marks: strong trans enh (highest expression), trans, and trans wk (transcribed but at different expression levels depending on the presence of H3K27ac H3K4me1, and H3K36m3), ZNF (Zinc Finger-related genes), poised, PC repr wk (weak Polycomb-mediated repression), bivalent (high level of repressive mark H3K27me3, and moderate levels of H3K4me1 and H3K4me3), het (genes associated with heterochromatin), PC repr (Polycomb-mediated repressed genes).

These metabolic reconstructions span 58 tissues and cell types. Blood cells account for the largest fraction of these models, with 356 T cell, 303 monocyte, 224 neutrophil, and 107 B lymphocyte lineage models **(Figure 1, left bar plot).** The median number of model reactions per cell and tissue type ranged from 2,360 (T cell) to 4,252 (liver) **(Figure 2A)**. The twelve model groups with the lowest median number of reactions are models for blood and immune cells, notably T cell, B lymphocytes, neutrophil, venous blood, and natural killer cell models **(Figure 2A)**. By contrast, hepatocyte, trophoblast, placenta, colon, and kidney models have the highest number of reactions, an indication of their greater metabolic complexity. The substantial size of the liver and hepatocyte models is consistent with the large number of liver-specific metabolic tasks such as drug metabolism, bile acid synthesis, homeostasis of carbohydrates, amino acids, lipids, and fat-soluble vitamins ^22,23^.

**Figure 2:**
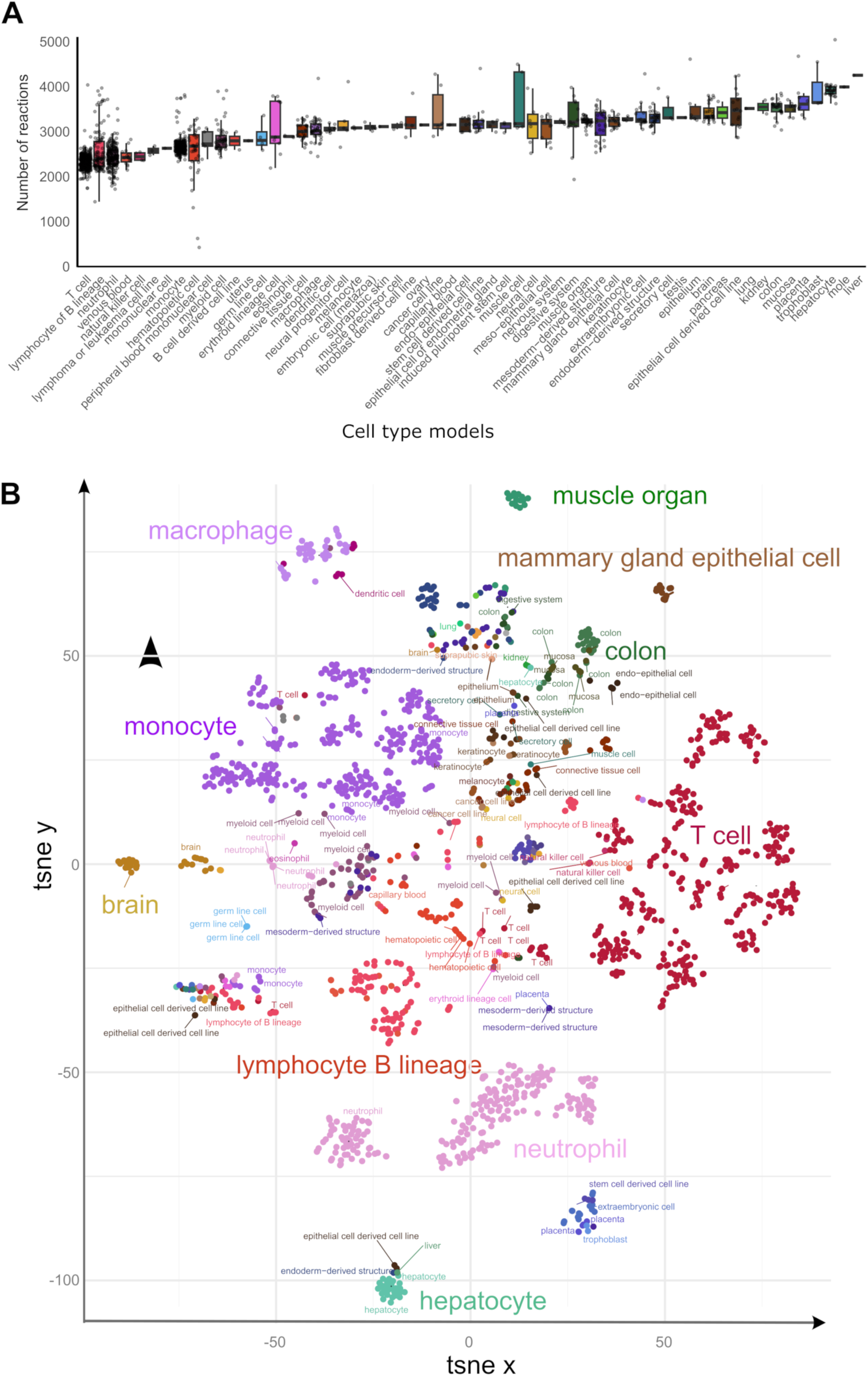
Sample models capture metabolic complexity across tissues and cell types. A) The distribution of the number of model reactions per cell type and tissue is represented as a box plot. The number of reactions in each model indicates the complexity of the metabolism of cell types and tissues. Error bars represent the upper and lower quartiles. Outliers are data points outside the range defined by Q1 - 1.5 interquartile range (IQR) and Q3 + 1.5 IQR. B) t-SNE embedding of sample models separates tissues and cell types. Sub-clusters are visible for models of some cell types such as T cells, neutrophils, lymphocytes of B lineage, and monocytes.

While many basic metabolic processes are shared across tissues and cell types, we hypothesized that individual tissues and cell types (1) fulfil core metabolic processes through tissue-specific reactions and regulatory mechanisms and (2) have tissue-specific metabolic processes for tissue-specific functions (*e.g.*, bile acid synthesis in hepatocytes). In support of these hypotheses, we observed that a t-SNE embedding of samples by metabolic models distinguished cell-type clusters (**Figure 2B**). This result is further supported by greater Jaccard similarity (in reaction composition) within tissue types than between tissue types **(Supplementary Figure S1A**). Notably, hepatocytes and neutrophils define the extremes of the first principal component (**Supplementary Figure S1B**), suggesting a highly distinct metabolism from other cell types. Taken together, these data demonstrate that metabolic models capture the cell type-specific metabolic character of cell types.

We next defined the set of ‘core’ metabolic reactions as the set of 541 reactions (from 10,600 total reactions) in Recon3D ^24^ which were found in over 99% of models. These include reactions in pathways required for cellular homeostasis, including reactive oxygen species (ROS) detoxification, fatty acid synthesis, squalene and cholesterol synthesis, purine synthesis, and nucleotide interconversion. Moreover, pathways implicated in the maintenance of the extracellular matrix, such as chondroitin sulphate metabolism, and energy metabolism, were present ubiquitously across all cell types and tissues **(Figure 3A)**.

**Figure 3:**
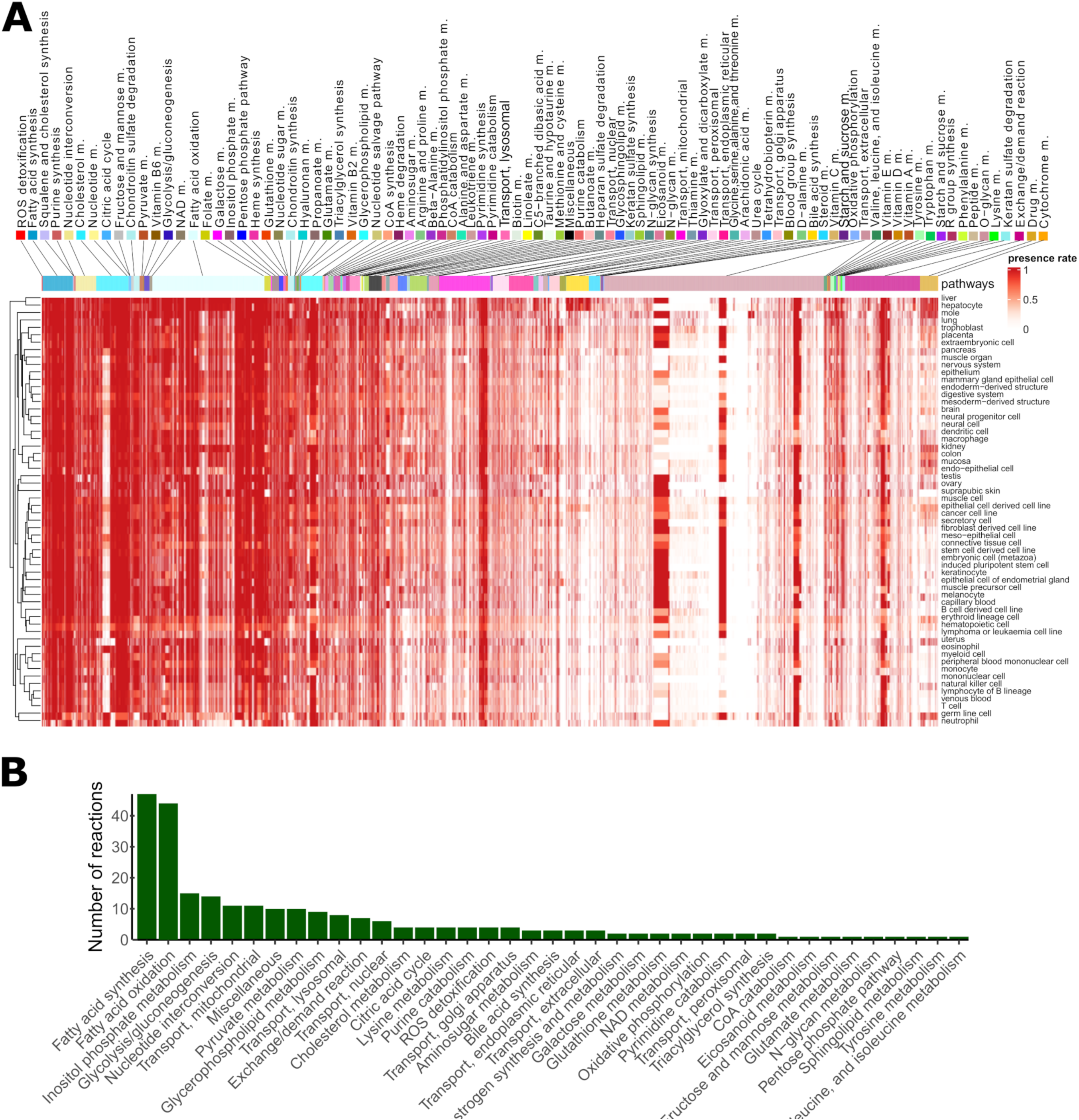
Metabolic models capture core and tissue-specific metabolic reaction pathways. **A)** Presence rates computed for each reaction and tissue type. Pathways were ranked according to their mean presence rate across models. Columns represent the reactions included in at least one of the models and rows represent the cell and tissue type. Some pathways (*e.g.*, ROS, fatty acid synthesis and squalene pathways) were included in most metabolic models, while other pathways (*e.g.*, drug metabolism, bile acid synthesis) were unique to hepatocytes. Even among the pathways with the highest presence rate, some reactions have a cell type-specific or sample- specific expression pattern. We observe a similar pattern for unique pathways (*e.g.*, bile, drug metabolism) which contain besides unique reactions a small fraction of ubiquitous reactions. B) The number of reactions per pathway present in at least 90% of the tissue and cell type models. Reactions of central carbon metabolism are among the reactions with the highest presence rate.

Next, we computed the number of reactions present in at least 90% of samples in each tissue or cell type. 253 reactions fulfil this criterion include reactions in fatty acid synthesis and oxidation, inositol phosphate metabolism, and glycolysis/gluconeogenesis. Notably, even for pathways commonly shared across all tissues and cell types, some reactions were specifically included only in some models (**Figure 3B and Supplementary Figure S2).** In contrast to these highly common reactions, 4,799 reactions were included in less than 1% of models (3,944 were never included, comprising redundant exchange and transporter reactions, drug metabolism reactions, and reactions not under gene control and thus not supported by the data) **(Supplementary Table S3)**. Some bile acid synthesis, drug metabolism, cytochrome C, steroid metabolism, and fatty acid oxidation pathways were unique to a single tissue or cell type, namely hepatocytes. However, we observed that other reactions in these pathways were present also in other tissues, suggesting that pathway activation or repression may occur at specific key reactions rather than at the level of the whole pathway.

Lymphoma or leukaemia cell lines, as well as blood and immune cells, have a more compact metabolism (*i.e.*, smaller number of reactions) **(Figure 3A)**, but also tend to have a lower reaction presence rate in most pathways, as evidenced by the brighter horizontal lines in **Figure 3A**. This finding suggests that, for some samples, these reactions with a lower presence rate can be shut down, possibly due to environmental signals or, in case of cancer cells, due to mutations. Some branches of fatty acid oxidation (FAO) were absent or had low presence rates in most cell types except for liver, placenta, trophoblast, extraembryonic cell, and mole. Finally, the reaction presence rate of metabolic models correlates with the median expression levels associated with the metabolic genes mapped to Recon3D, demonstrating the sensitivity of model building pipeline **(Supplementary Figure S3)**. Taken together, the reconstructed metabolic models reveal core and tissue-specific metabolic characteristics.

### Core and unique metabolic genes are associated with higher enhancer activity and expression levels than non-housekeeping genes shared by multiple tissues

While genes controlling core metabolism, by definition, must be expressed ubiquitously, we sought to determine whether these commonly expressed genes are differentially regulated across cell types. We defined a set of core genes (n=3,530) as genes with active ChromGene ^20^ annotation across all EpiATLAS cell types and tissues (Methods). We also defined a set of core metabolic genes (n=306) as the subset of core genes present in Recon3D. Similarly, we defined unique (n=1,013) and unique metabolic genes (n=121) as the set of genes active in only a single tissue or cell type. The core gene set was enriched for genes implicated in the metabolism of RNA and processing of introns, while core metabolic genes were enriched in mRNA splicing, translation, energy production, biosynthesis pathways, lipid metabolism, and ROS metabolism **(Supplementary Table S3)**. Both core gene sets were strongly enriched for common essential genes obtained within the DepMap project ^25^ **(Figure 4A)**. 934 out of 1,549 common essential genes and 48 out of 79 common essential metabolic genes were core and core metabolic genes, respectively. Core genes and core metabolic genes together captured 60% of the common essential genes. The missing 31 essential metabolic genes were associated with active ChromGene states (1-6) in most tissues and cell types, with one exception, gamma-glutamyl transferase (*GGTLC2*), which carried a repressive chromatin state across most samples (**Supplementary Figure S4)**. We observed similar patterns for non-metabolic core essential genes: except for a few genes that are associated with a repressive state in most cell types, core essential genes carry activating states (ChromGene state 1-6) in most cell types (**Supplementary Figure S4)**. The enrichment for overlaps between common essential and core genes implicates these genes in key metabolic pathways that are highly conserved across cell types and essential for cell survival.

**Figure 4:**
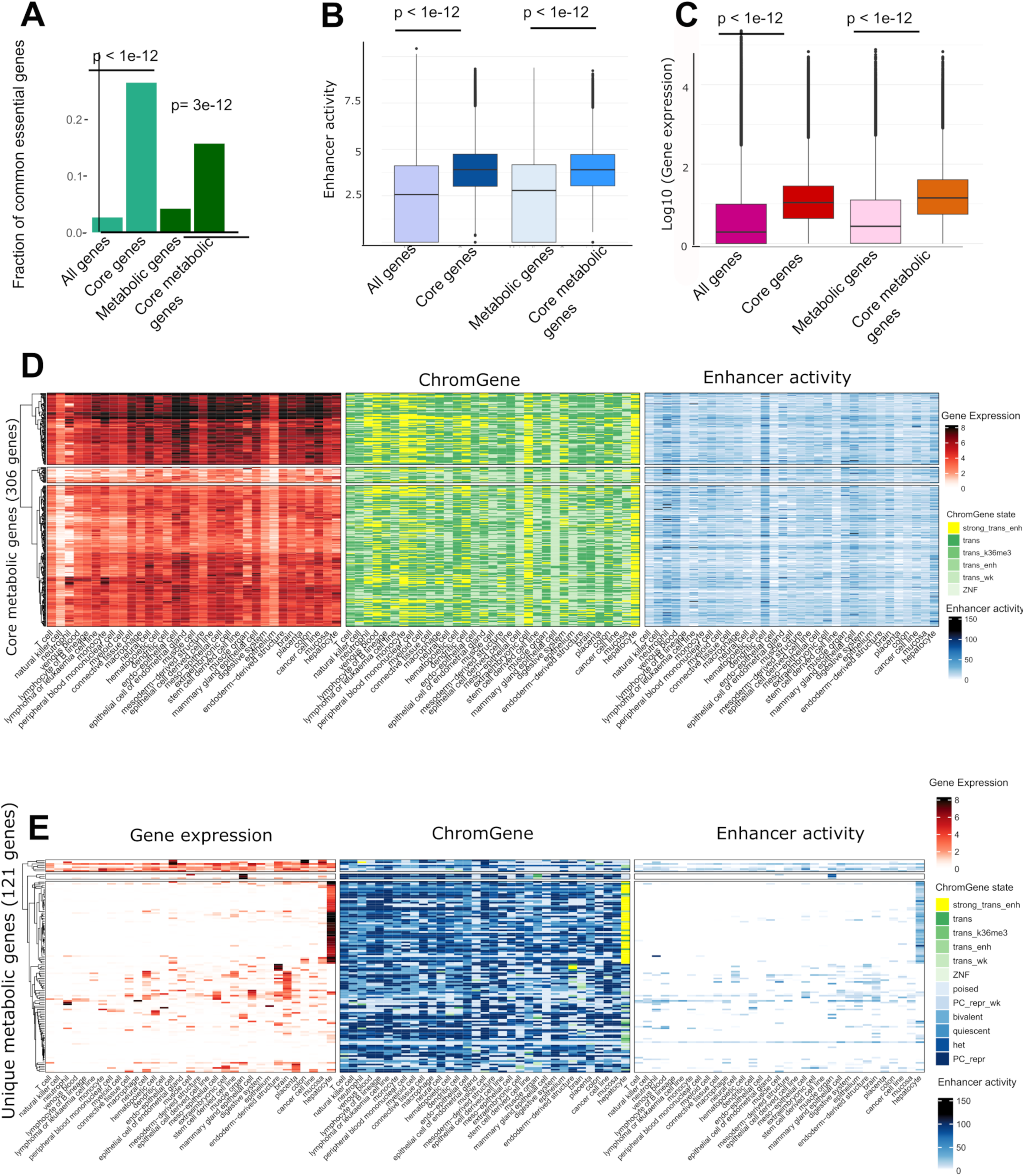
Core and core metabolic genes are essential and exhibit high enhancer activity and high expression. **A)** Core and core metabolic genes are enriched for essential genes common to most cell types (hypergeometric p-value < 3e-12). **B)** Core and core metabolic genes have higher enhancer activity scores than genes on average (t-test p-value <1e-12) and **C)** higher median expression values (t-test p-value <1e-12). **D)** Core metabolic genes have high expression values (log2 TPM, left), are linked to active ChromGene states (middle), and have a high enhancer activity across most cell types and tissues (right). Rows contain the 306 core metabolic genes and columns contain the cell types with at least 3 samples for each data type. **E)** Similar to D but representing 121 unique metabolic genes. Unique genes have a high expression in a single tissue or cell type and have low or no expression in the other tissues. There is also a high number of Polycomb-repressed genes, heterochromatin states, and low enhancer activity scores for these unique genes in other tissues.

We next used the generalized Activity-By-Contact (ABC) method ^17,26^ to examine the enhancers linked to core and core metabolic genes. For each gene, we computed an enhancer activity score by summing the activity of all enhancers predicted to interact with a gene per cell type or tissue (Methods). Consistently, higher median enhancer activity scores were observed for the core and core metabolic genes **(Figure 4B)**. Also, the increase of the median gene expression was more pronounced for the core and core metabolic genes compared to all genes and all metabolic genes **(Figure 4C).** This is consistent with the previous observation that core fitness genes have substantially higher expression than non-fitness genes ^27,28^. Also, core metabolic genes are associated, by definition, with activating ChromGene states 1-6 and a higher enhancer activity **(Figure 4D)**. There was a substantial number of genes with ChromGene state 1 (corresponding to highly expressed genes with strong signals of active marks such as H3K4me1 and H3K27ac) among the genes expressed in hepatocytes and extraembryonic cells, as well as neutrophils, monocytes, lymphocyte B lineage, venous blood, peripheral blood mononuclear cells, and myeloid cells.

For unique genes, we found both higher expression and higher enhancer activity to correspond to an active ChromGene state **(Figure 4E, Supplementary** Figure 5). Of the 1,013 unique genes, 177, 144, and 109 were specific to hepatocytes, brain, and muscle, respectively **(Supplementary Figure S6A)**. These sets were enriched for gene ontology terms matching their cellular identities (*e.g*., brain enriched for protein-protein interactions at synapses). Among the 121 unique metabolic genes, as many as 67 were hepatocyte-specific genes and were consistently enriched for tasks and pathways specific to hepatocytes **(Figure 4E, Supplementary Figure S6B, and Table S4-S5)**. The high expression of these unique gene sets matched high enhancer activity scores while low expression and enhancer activity was observed in other tissues. Polycomb-mediated repression was the most common repressive chromatin state at the cell type-specific genes **(Figure 4E)**. However, the number of Polycomb-repressed genes among the 121 unique genes varied between tissues. Muscle cell, lymphoma and leukaemia samples, and cancer cell lines had 62, 60 and 57 Polycomb-repressed unique metabolic genes, respectively. This suggests that besides gene activation in unique cell types, repressive states, and extensive Polycomb-mediated repression, could also contribute to shape metabolic identity.

### TFs controlling metabolic identity show cell type-specific activity

Next, we asked whether the high and consistent expression of core genes and core metabolic genes was controlled by the same TFs across tissues, or whether core metabolism is subject to cell type-specific regulation. To this end, we integrated cell type-specific enhancer-gene interactions to obtain sets of regulatory elements linked to each core and core metabolic gene for each cell type **(Figure 5A)**. Many TFs were uniformly enriched at enhancers associated with core genes across different cell types **(Figure 5A).** Conversely, we observed that some TF regulators of core metabolic genes were uniquely enriched in specific cell types and tissues. Overall, 264 non-redundant TF motif modules and tissue-specific regulatory regions were significantly enriched at enhancers associated with core metabolic genes (FDR<0.01), and 41 of these had absolute enrichment scores over 1.3. These TFs control common processes such as growth, proliferation and differentiation (EGR)^29^, oncogenic and embryonic development (TFAP2, SPDEF, ZFX, ZIC1, ZBTB7A, PAX, GLIS) ^30,31,40,32–39^, regulation of chromatin structure (MECP2, HINFP, MBD2, CTCF, KAISO)^41–46^, and cell cycle (E2F, RUNX)^47,48^. RUNX1 is expressed in hematopoietic cells ^49^, but also plays a role in the innate immune system of non-hematopoietic cells ^50^. Nuclear receptors broadly regulate metabolism and inflammation ^51^. HINFP and GMEB2 showed the largest variation in enrichment.

**Figure 5:**
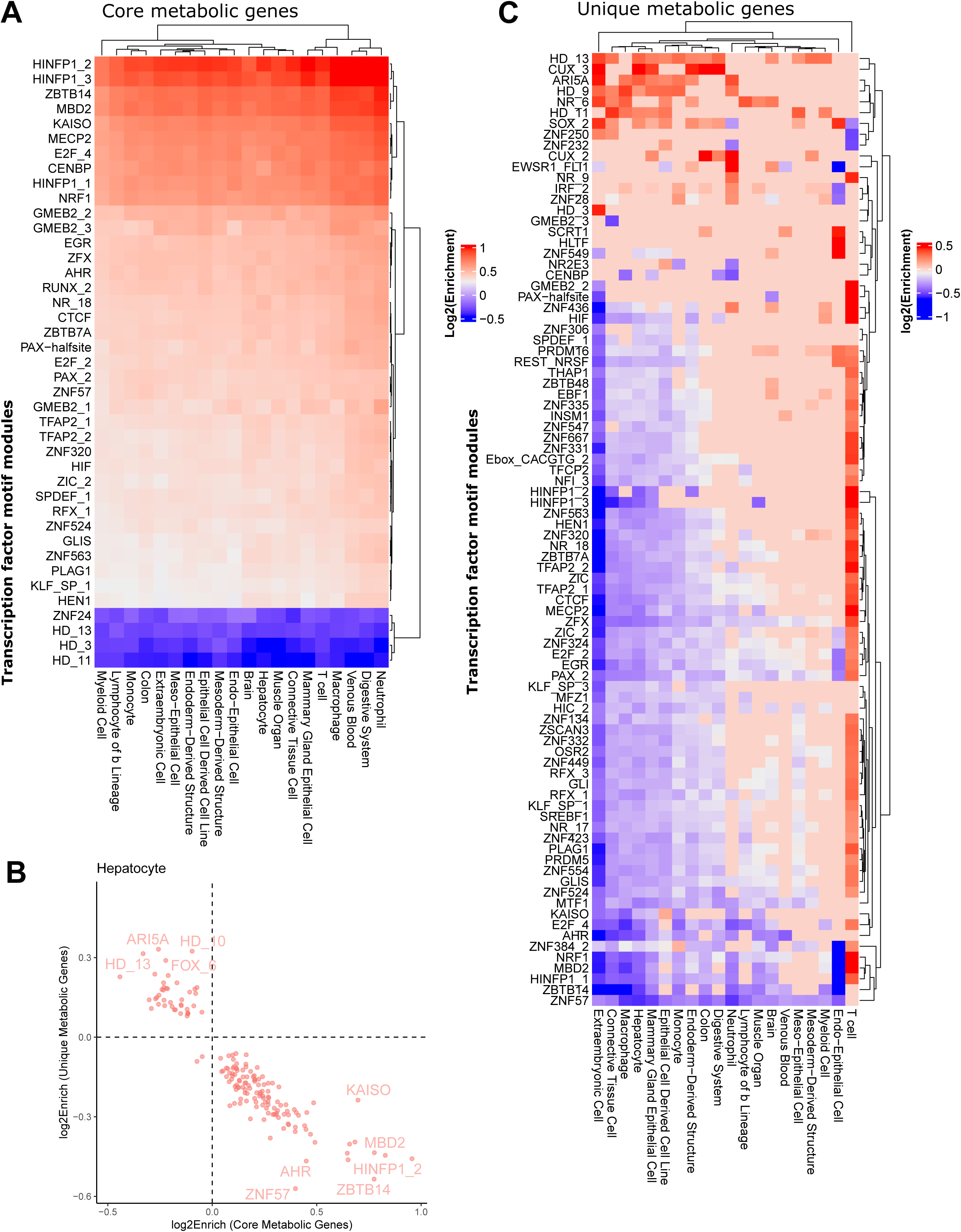
**Core metabolic genes and unique metabolic genes are enriched for different TF sets.** A) The enhancer-gene interactions were used to obtain a union set of regulatory elements for ten samples randomly chosen from the cell types and tissues with more than 10 samples for the core and core metabolic genes. TFs implicated in common biological processes such as growth proliferation, and development response to stress or environmental changes were enriched for core metabolic genes. Only TFs with absolute enrichment above 1.3 are depicted. B) Negative correlation between enrichment (log2, FDR<0.01) of TFs for core metabolic genes vs unique metabolic genes. C) Same analysis as A) but for unique genes. Only TFs with an absolute enrichment above 1.3 are depicted.

Interestingly, we observed an inverse association between TF enrichment values for core metabolic genes and unique metabolic genes **(Figure 5B)**, suggesting distinct upstream regulators of core and tissue-specific metabolic processes. Consistently, some TF modules with negative enrichment values in core metabolic genes, such as HD-3 and HD-13, have high enrichment values for the unique metabolic genes **(Figure 5C)**.

Other TF groups with high enrichment at enhancers of unique metabolic genes included CUX^52^ (implicated in cell cycle control and DNA repair), and SOX (SOX9 plays a role for example in liver homeostasis and regeneration^53^).

Most of these TFs and motifs have a wide number of target genes and were shown to be dysregulated in numerous cancers ^54–56^. A more detailed understanding of the cell type-specific control of core and unique genes could allow targeted interventions in specific cell types such as diseased cells.

### Epigenetic control shapes metabolic identity by regulating solute carrier genes in a tissue-dependent manner

Encouraged by our findings that unique genes are marked by Polycomb-mediated repression in other tissues, we next sought to determine if this repression was targeted to genes controlling key nodes in metabolic pathways or broadly across genes in metabolic pathways. In total, 750 metabolic genes were associated with Polycomb-mediated repression in at least one tissue **(Figure 6A, Supplementary Figure S8)**. We observed that leukaemia or lymphoma cell lines (409 metabolic genes), muscle cells (348), natural killer cells (286), and cancer cell lines (282) have the highest number of Polycomb-repressed metabolic genes. Conversely, stem cell-derived cell lines, mammary gland epithelial cell lines, and brain samples have the lowest number (38, 55 and 67). Mammary gland epithelial cells are instead enriched for poised promoters **(Supplementary Figure S9),** possibly linked to their precise, highly regulated response to hormonal signals. The poised and Polycomb-repressed ChromGene states are mutually exclusive, explaining the lower number of Polycomb-repressed genes in the cells carrying many poised promoters. The 409 Polycomb-repressed genes in leukaemia or lymphoma cell lines are enriched for pathways such as bile secretion, purine metabolism, and steroid hormone biosynthesis, which are mostly active in hepatocytes. Furthermore, in all the tissue types Polycomb-repressed genes are enriched for genes encoding for solute carriers (SLCs) **(Figure 6B)**. The proportion of SLCs among the Polycomb-repressed genes is between 15.8% (1.3-fold enriched) and 25.5% (2.1- fold enriched) for stem-cell derived cell lines and epithelial cells of the endometrial gland, while overall they represent only 12.4% of the metabolic genes **(Figure 6A)**. For lymphoma or leukaemia cell lines, 76 SLCs (out of a total of 229 SLCs, p = 1.e-5) and 8 (ATP)-binding cassette transporters (out of 22, p = 0.08) **(Figure 6A)** are Polycomb-repressed.

**Figure 6:**
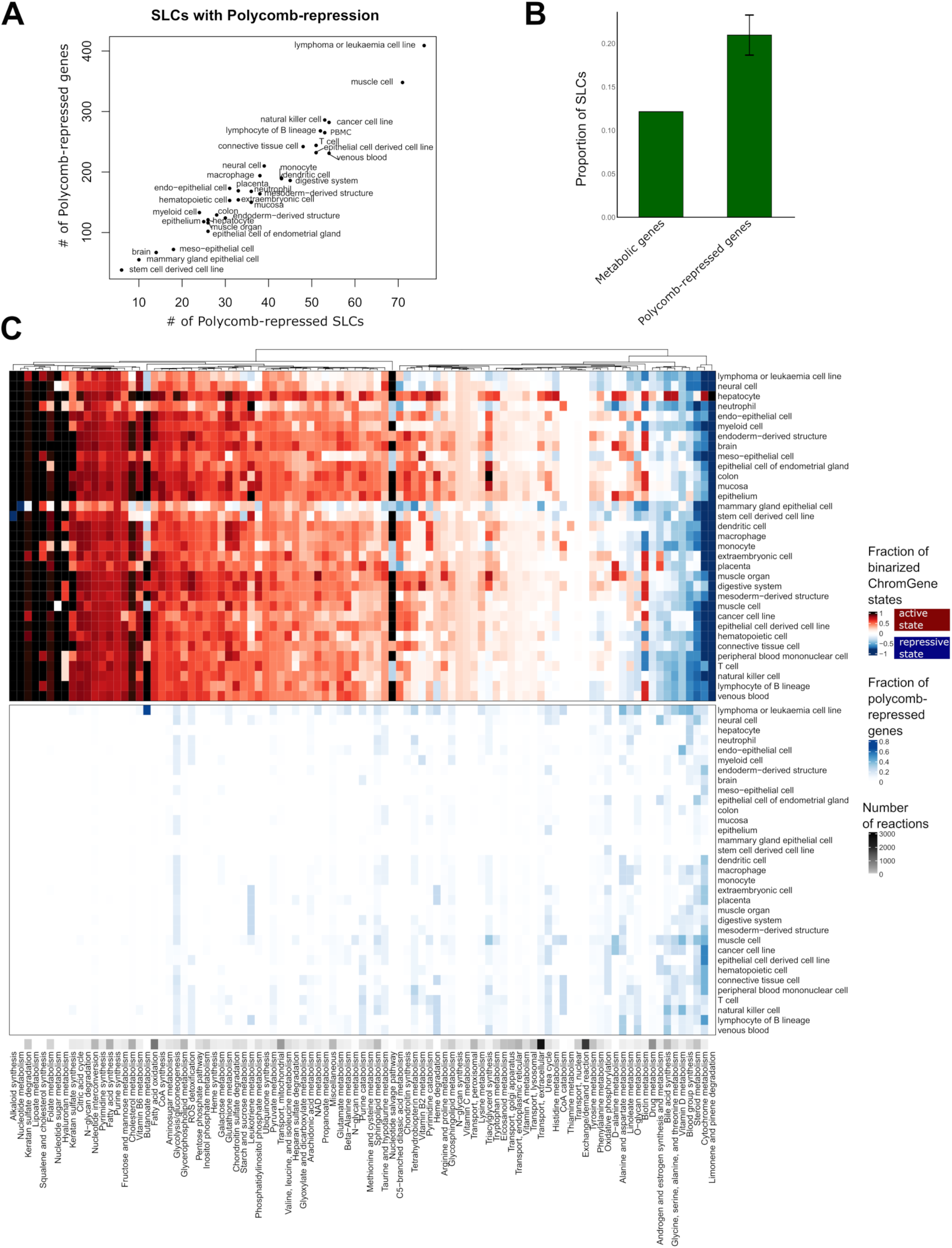
Polycomb repression shapes metabolic networks by repressing solute carriers. A) Number of Polycomb-repressed SLCs over number of Polycomb-repressed genes across cell types. B) Proportion of SLCs in metabolic genes and Polycomb-repressed genes. C) Presence rate of different ChromGene states for the different metabolic pathways. Top: ChromGene states were binarized for visualization purposes. Active ChromGene states 1-6 were associated with a score of 1, and inactive ChromGene states between 7 and 12 with a score of –1. Gene expression was then mapped using the Gene-Protein-Reaction rules of the metabolic network to obtain a score for each reaction. A mean score for each pathway was computed by summing the individual scores and dividing by the number of reactions (top). Bottom: Same plot but considering ChromGene state 12 (Polycomb-repressed, bottom panel). The black ribbon depicts the number of reactions per pathway.

To analyse the distribution of Polycomb-mediated repression across metabolic pathways, we compared the binarized ChromGene states and the Polycomb-repression distribution (ChromGene state 12). A core metabolism cluster highlighting mostly the same pathways as in **Figure 3A** is enriched for active chromatin state **(Figure 6C top)** across all tissues and cell types and depleted of any Polycomb-repression (**Figure 6C bottom**). Several pathways have a presence rate of above 0.6 for ChromGene state 1 in three or more tissues **(Supplementary Table 8**, **Supplementary** Figure 10). Hepatocytes have a presence rate above 0.6 for 21 pathways indicating once again high metabolic complexity. For Polycomb-mediated repression only a few pathways have a presence rate above 0.3 in more than one cell type (**Supplementary Figure S10).** Leukaemia and lymphoma cell lines had the highest presence rates for Polycomb-repression in butonate metabolism, blood group synthesis, alanine and aspartate metabolism, vitamin D, O- glycan metabolism, glycine, serine, alanine, and threonine metabolism. Unlike what is seen for active chromatin states, Polycomb-repression never affects every reaction of a pathway, but only some specific branches of pathways (**Supplementary Table S9**).

### Transporters and other pathway entry points are key nodes for epigenetic regulation of the metabolic networks

As SLCs are enriched for Polycomb-mediated repression, and hence could serve as switches within the metabolic network, we tested whether SLCs were also enriched among the uniquely expressed metabolic genes **(Figure 7A)**. Within the 121 unique genes, we observed an enrichment for cytochrome P450 (CYP) enzymes (12 out of 48, p = 3e-05), which is consistent with the enrichment for hepatocyte-specific genes. Furthermore, 33 out of 371 genes controlling transport reactions, were among the 121 unique metabolic genes (p = 0.02), (3 ATP binding transporters, 21 SLCs and the transporter genes NPC1L1, SULT2A1, AQP8, PRODH2, RBP2, RBP4, RHCG, ATP1A3, ATP1A4).

**Figure 7:**
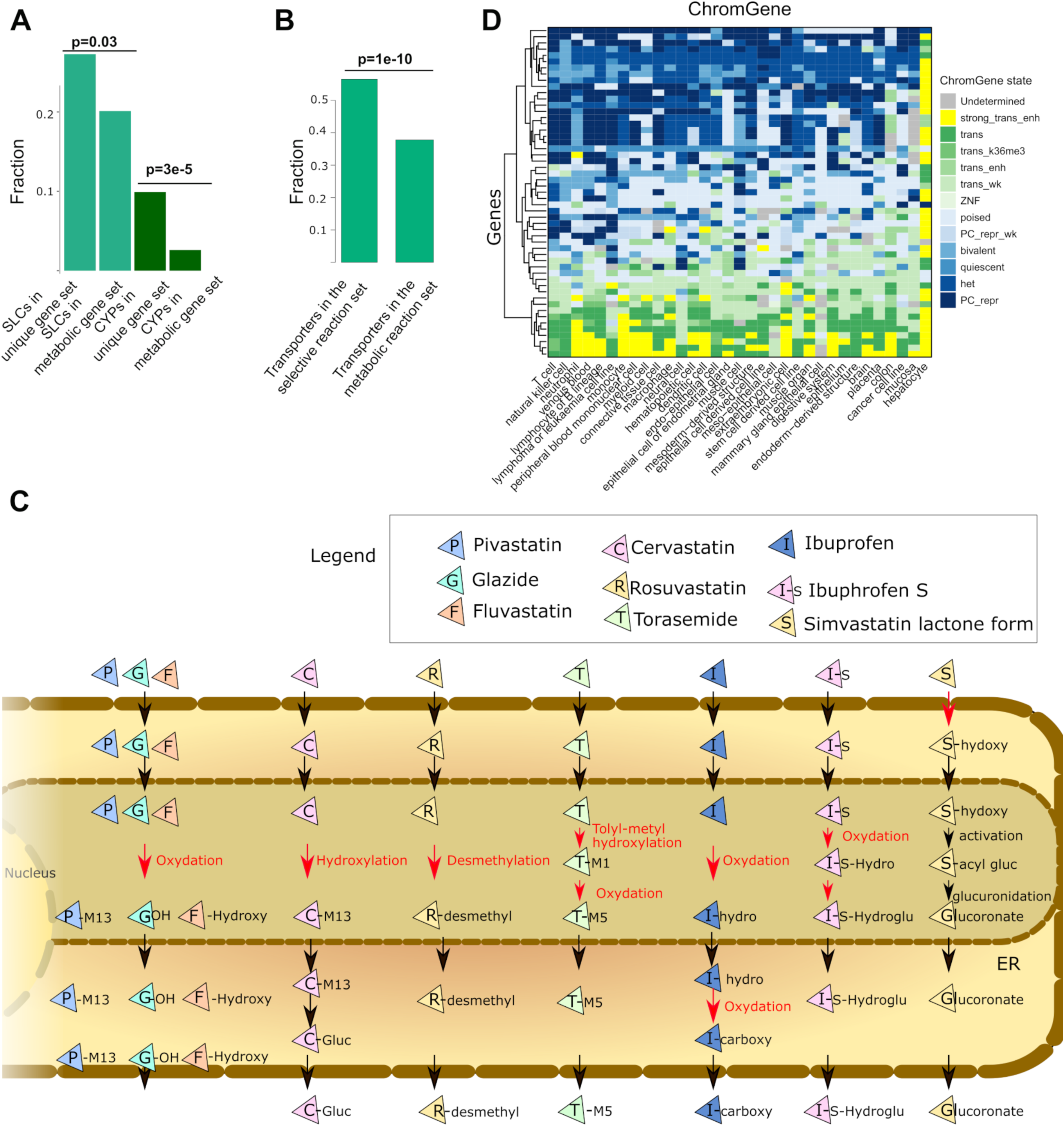
Transporter and CYP genes are enriched among unique genes; transport reactions are enriched among unique reactions. A) Among the 121 unique genes, 12 out of 48 genes and 33 out of 271 genes code for CYP proteins and transporter coding genes, respectively (hypergeometric test). B) Among the 256 unique reactions, 152 out of 2,241 are transporter reactions (hypergeometric test). C) Schematic representation of the drug metabolism pathway in hepatocytes. Reactions under unique control are depicted in red. These unique reactions are often located in the endoplasmic reticulum and generally after a transport reaction between two compartments or an import into the cell. D) ChromGene states linked to the bile acid genes are depicted. Some genes implicated in drug metabolism are often uniquely expressed in hepatocytes and control 13 out of 573 reactions. In other tissues, these genes are often Polycomb repressed.

To take advantage of the structural information of the metabolic reconstruction, the 121 uniquely active metabolic genes were mapped to the networks to unravel key switches. Among the 5,938 reactions under known gene control in the generic Recon3D reconstruction, 3,744 reactions are associated with transport. Of these, 2,241 are controlled by gene products, while the remaining 1,503 are diffusion reactions.

Among the 269 unique reactions under a selective expression regulation (controlled by one or several of the 121 genes), 151 were transporter reactions controlled by 17 genes. Consequently, an enrichment for unique expression regulation for the transporters was also found at the reaction level (p = 1e-10) **(Figure 7B)**. 97, 28, 6, and 4 out of the 151 transporter reactions are controlled by SLCO1B1, SLC22A10, SLC5A8 and SLC22A2, respectively. Besides, extracellular transporters (151/2652), bile acid synthesis (27/185), drug metabolism (13/573), glycine, serine, alanine, and threonine metabolism (11/47), cytochrome C (8/15) and glycolysis/gluconeogenesis (6/42) have the largest number of reactions controlled by unique genes. These reactions are often importing metabolites into the cell or are the first reaction after a transport reaction to the endoplasmic reticulum or another compartment, as exemplified by the drug metabolism pathway **(Figure 7C)**. Accordingly, drug metabolism is inactivated in other tissues by repressing their import in the cell or repressing enzymes in the endoplasmic reticulum metabolising drugs **(Figure 7C)**. Similarly, we have already shown the importance of regulation of entry points for the bile acid pathway (IHEC flagship paper, under preparation). This pathway has a large fraction of genes and associated reactions with a ChromGene state of 1 (highly expressed in hepatocytes) **(Figure 7D)**. In other tissues, this pathway seems to be mostly inactive due to the Polycomb-repression of genes controlling switches in the network **(Figure 7D)**. Overall, the metabolic network analysis shows an orchestrated epigenetic regulation combining activating and repressing signals to control key nodes in the networks in a cell-type-specific manner.

## Discussion

In this study, we reconstructed 1,555 metabolic network models capturing core metabolic pathways and processes and the basis of the distinct metabolic identities across cell types. Both core and unique metabolic genes have a higher enhancer activity, and consequently, higher expression values, compared to the remaining metabolic genes, providing robustness to the control of vital processes, consistent with the high number of common essential genes among the core genes. 60% of common essential genes were included among the core genes. We defined core genes based on strict criteria (active ChromGene states for all cell types and tissues), and consequently, many of the common essential genes missing from the core gene set were associated with active chromatin states in most cell types. Gene set enrichment analysis further showed that unique genes (considering both metabolic and non-metabolic unique genes) are sufficient to infer the identity of the tissue and cell types. Unique genes exhibit active ChromGene state in the cell type they are expressed in and strong Polycomb repression in other tissues. Notably, stem cells exhibit fewer Polycomb-mediated repressed genes, suggesting a relationship between differentiation stage and repressive mechanisms. Conversely, we observe more Polycomb-repressed genes in cancer cell lines, consistent with Polycomb dysregulation in cancer ^57^.

We observed that transport reactions, specifically SLCs, were enriched for unique metabolic genes and often Polycomb-repressed. This is consistent with the strict regulation of transporters, as SLCs control the import of metabolites, and channel them into storage or degradation compartments. While precise transporter regulation is energetically expensive, it enables more robust regulatory systems where regulation is enforced only at key points of the networks. SLCs have diverse expression patterns with potential substrates that range from ions, lipids, vitamins, waste products to drugs ^58,59^. Some members of the SLC superfamily are ubiquitously expressed, while others are cell- or even context-specific. Mutations in 20 % of SLC genes are linked to Mendelian ^60^ diseases. SLCs are also often dysregulated in cancer, as the ability to control non-vital metabolic pathways can facilitate tumor growth. SLCs are often hypermethylated in cancer, and SLC expression has been linked to cancer clinical outcomes ^61^.

Finally, we showed that core genes are under the regulatory control of TFs linked to essential biological processes such as growth, proliferation, and chromatin structure. However, most of these TFs are not essential, suggesting redundant regulatory systems. Notably, TFs controlling core metabolism were often depleted in the unique gene sets. As an example, binding sites for KAISO were enriched in core metabolism and depleted in the unique gene sets of several tissues. KAISO regulates development in normal tissue and plays a role in tumorigenesis through its impact on cell cycle and methylation. Similarly, binding sites for HINFP, which is critical for late embryogenesis cell cycle control ^62^, were broadly depleted from unique gene sets despite strong enrichments in the core metabolic set. HINFP is crucial in the control of the cell cycle during late embryogenesis ^62^.

## Conclusion

In summary, our analysis, empowered by the comprehensive EpiATLAS resource, has characterized the metabolic identities across 58 cell types and tissues, delineated core and tissue-unique metabolic components, and, by integrating cell-type-specific gene-level chromatin states and enhancer-gene interactions, identified regulatory elements, upstream transcription factor regulators, and key pathway nodes controlling metabolism. We observed that transporters, specifically SLCs, and first reactions of pathways were enriched for high expression, active chromatin state, and Polycomb-mediated repression. This analysis serves as the foundation for uncovering regulatory sites that govern metabolic identity in human cells, providing a basis for tissue-specific metabolic reprogramming.

## Methods

We accessed RNA-seq profiles (1,555 samples across 58 cell types and tissues), ChromGene HMM states (1,698 samples), and enhancer activity data (1565 samples) from the IHEC EpiATLAS resource (https://epigenomesportal.ca/ihec/, International Human Epigenome Consortium, EpiATLAS - a reference for human epigenomic research, in preparation) ^16^. Of these, 965 samples had expression, ChromGene, and enhancer activity data and 938 samples belonged to tissues and cell types with at least 3 samples. RNA-seq and ChromGene data were accessed from the IHEC sftp server in batch mode. The summed enhancer activity on gene-level was taken from the IHEC sftp as well (‘gABC_GeneSummedEnhancer_geq2.txt.gz’). This file sums the H3K27ac signal of all enhancers of a gene, based on interactions predicted with the gABC-score^17^. The score of genes without sufficient activity at any of their promoter (≥2 H3K27ac signal ±200 bp around any annotated TSS) was set to zero.

### Reconstruction of metabolic models

1,555 sample models were reconstructed for each RNA-seq sample by the rFASTCORMICS ^5^ using Recon 3D ^24^ as input reconstruction, as previously described. Briefly, rFASTCORMICS discretizes RNA-seq values into expressed, not expressed, and undetermined expression status. Expressed and not expressed genes are mapped to the generic reconstruction (here Recon3D) to form two sets of reactions: core and inactive reactions. Core reactions are supported by the data and should be included in the model. Inactive reactions are under the control of genes that are not expressed and hence are removed from the network. rFASTCORMICS uses FASTCORE^9^ to build a flux-consistent network by adding to the set of core reactions (which are not blocked due to the presence of inactive reactions in the same pathway), a next-to-minimal set of non-core reactions. The models were not medium-constrained.

### Jaccard Similarity Index, t-SNE, and Principal Component Analysis

For each pair of models, the Jaccard Similarity Index (JSI) was computed, and the models were then clustered according to the JSI using the Euclidean distance. To verify that the clustering on the models captured metabolic variations, we performed principal component analysis on the metabolic genes retrieved from Recon3D using the *factoextra* package ^63^.

We generated a t-SNE embedding of the sample models using the *tSNE* package^64^ in R. Briefly, we constructed a binary matrix encoding reaction presence across 10,600 Recon 3D reactions in 1,555 samples. We excluded reactions absent from all samples in our dimensionality reduction analysis.

### Core, core metabolic, unique, and unique metabolic genes

A consensus ChromGene score is the most represented cell type and tissue gene mark. If the most represented mark accounts for less than 50% of the samples and is an activating mark and the other marks are repressive then a score of zero is assigned. Similarly, if the majority mark accounts for less than 50% of the samples and is repressive and the other marks are activating marks, the samples get a consensus score of zero. We defined a set of core genes as those with active consensus ChromGene states (states below 7) for all 33 cell types in the set of 938 EpiATLAS samples with expression profiles, ChromGene annotations, and enhancer activity data. Conversely, we defined unique genes as those which contained active consensus ChromGene states in only one tissue. We then defined core metabolic and unique metabolic gene sets as the subsets of core and metabolic genes included in Recon3D. The log2 expression values, consensus ChromGene states, as well as the enhancer activity were mapped to the metabolic genes of Recon3D and plotted with the ComplexHeatmap package ^65^.

### Enrichment of common essential genes among core genes and gene set enrichments

To validate the core sets, we computed the enrichment of common essential genes (genes essential across all cell lines in DepMap) downloaded from https://depmap.org/portal/download/all/

(CRISPRInferredCommonEssentials.csv) in the core sets using a hypergeometric test. Gene ontology enrichment terms for all four sets were downloaded from EnrichR ^66^ (https://maayanlab.cloud/Enrichr/).

### Presence rate of ChromGene states

The consensus ChromGene states were mapped to the metabolic genes, and a ChromGene score for each reaction was computed using the Gene-Protein-Reaction rules (GPR rules). GPR rules state which genes control a reaction and if the gene products act as protein complexes (denoted by BOOLEAN AND), or as isozymes (BOOLEAN OR) *e.g.*, Gene 1 AND Gene 2 OR Gene 3. To obtain a single score for each reaction, the maximum ChromGene value was taken for genes linked by a BOOLEAN OR and the minimum for BOOLEAN AND. The ChromGene was binarized to obtain a negative (−1) and a positive score (1) for ChromGene states below or equal to 7 and above 7, respectively. A mean score was then computed for each pathway. A similar approach was used for the genes linked to ChromGene 12 (Polycomb-repression) (Figure 6).

### Enrichment of transporter genes and reactions

Hypergeometric tests were performed to assess the enrichment of SLCs and CYPs in unique metabolic genes. Furthermore, genes controlling reactions belonging to the transporters subsystems (pathways) in Recon3D were also assessed for enrichment. Finally, reactions of the transporter reactions were also tested for enrichment among the set of unique reactions (reactions that have an activating consensus ChromGene in a unique cell type and tissue).

### Transcription factor motif enrichment analysis

To identify upstream regulators of shared and tissue-specific metabolic gene regulation, we performed overlap enrichment analysis between transcription factor motif occurrences and metabolic gene-linked regulatory regions. Briefly, we began with the previously described sets for core and unique metabolic genes. For a particular sample, we limited these sets to genes with active ChromGene states in the corresponding tissue type. We used the previously described set of sample-specific enhancer-gene interactions to identify sets of gene-associated regulatory elements. We used the union set of regulatory elements from 10 samples in each of 20 tissue categories in our analysis. We accessed the human reference genome-wide motif occurrences of 280 non-redundant archetypal transcription factor motif modules ^67^. We calculated a tissue-specific binomial-based enrichment between pathway-associated enhancers and motif archetype occurrences ^68^ using the joint set of all core and unique gene regulatory regions as background regions.

To identify tissue-specific regulators of core and unique metabolic processes, we visualized enrichments across tissue categories. We pruned the resulting heatmaps by retaining motif modules with absolute enrichment of at least 1.3, for modules regulating core genes and unique metabolic genes.

### Ethics approval and consent to participate

Not applicable.

### Consent for publication

Not applicable.

### Availability of data and materials

The transcriptomic and epigenomic data supporting the conclusions of this article are available in the IHEC EpiATLAS resource repository, https://epigenomesportal.ca/ihec/, International Human

Epigenome Consortium, EpiATLAS - a reference for human epigenomic research, in preparation) ^16^.

The reconstructed metabolic models can be obtained at https://github.com/sysbiolux/IHEC_metabolism/.

## Funding

This research was funded in part by the Luxembourg National Research Fund (FNR), grant reference [PRIDE15/10675146/CANBIO]. D.G. was supported by funding from Fondation du Pélican de Mie et Pierre Hippert-Faber. Research reported in this publication was supported by the National Institute of General Medical Sciences of the National Institutes of Health under Award Number “T32 GM007748”. The content is solely the responsibility of the authors and does not necessarily represent the official views of the National Institutes of Health. For open access, the authors have applied a Creative Commons Attribution 4.0 International (CC BY 4.0) licence to any Author Accepted Manuscript version arising from this submission.

### Authors’ contributions

T.S., L.S., M.S., and M.K. conceived and managed the project. M.P.P., D.G., R.J.M., A.R.C., and D.H. performed all the analysis. All authors have read and approved the final manuscript.

## Acknowledgements

The computational experiments presented in this paper were carried out using the HPC facilities of the University of Luxembourg ^69^, https://hpc.uni.lu). All schematic representations were created with BioRender.com, R, or inkscape.

---

**Supplementary Figure S1:**
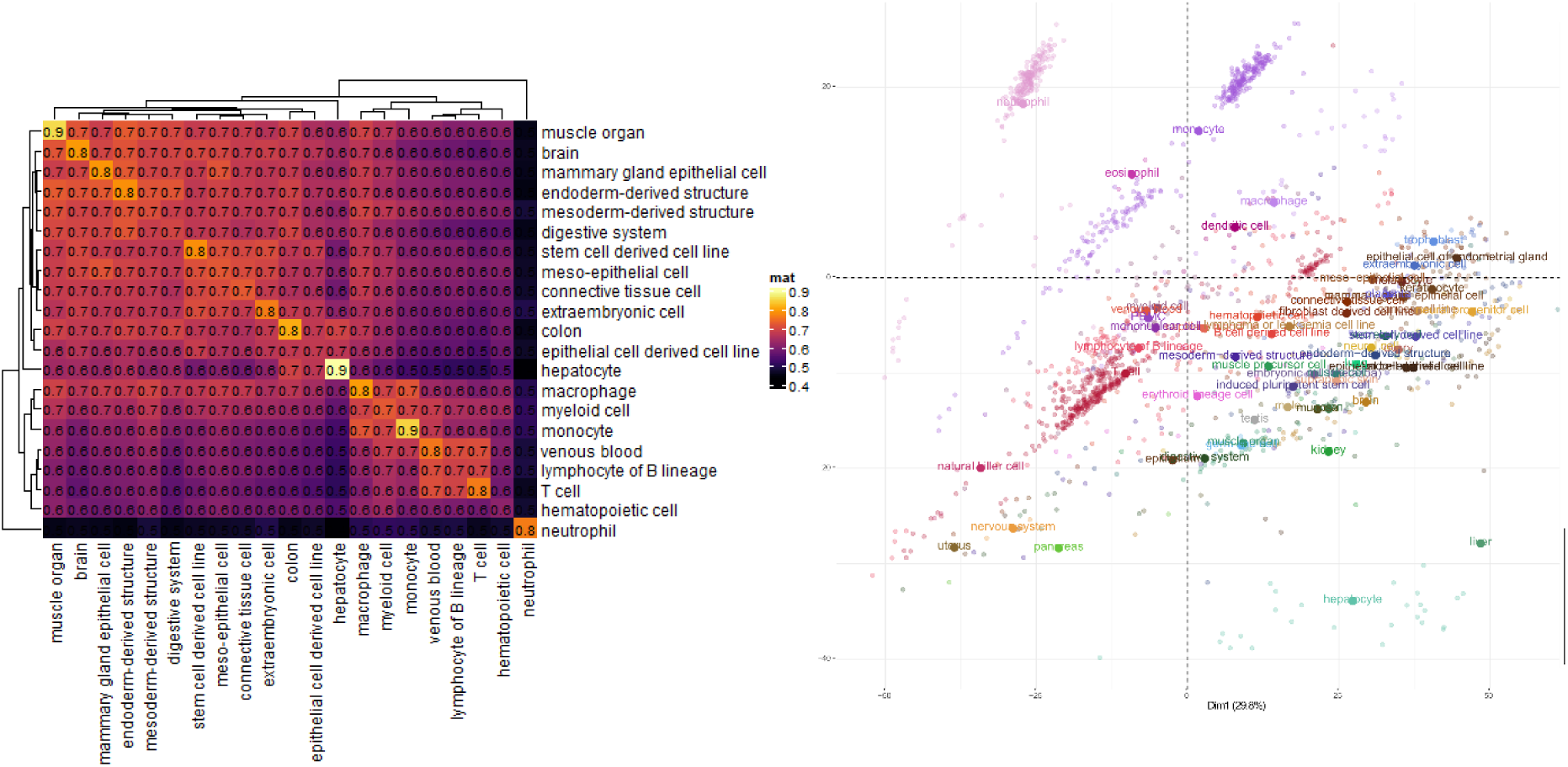
Metabolic models capture the metabolic identity of tissues and cell types. A) Hierarchical clustering of metabolic models by median Jaccard Similarity Indexes. A Jaccard Similarity Index (JSI) was computed for each pair of models. Then a median JSI was computed for each combination of cell types and tissues. The similarity within cell types and tissue tended to be larger than between different cell types and tissues. B) Principal component analysis of the metabolic gene expression data. Neutrophils are separated from the other blood cell samples. Non-blood cells are separated from the blood cells along PC1. The hepatocyte and liver samples like neutrophils also form a distinct cluster but at the other extremity of PC1 and PC2.

**Supplementary Figure S2:**
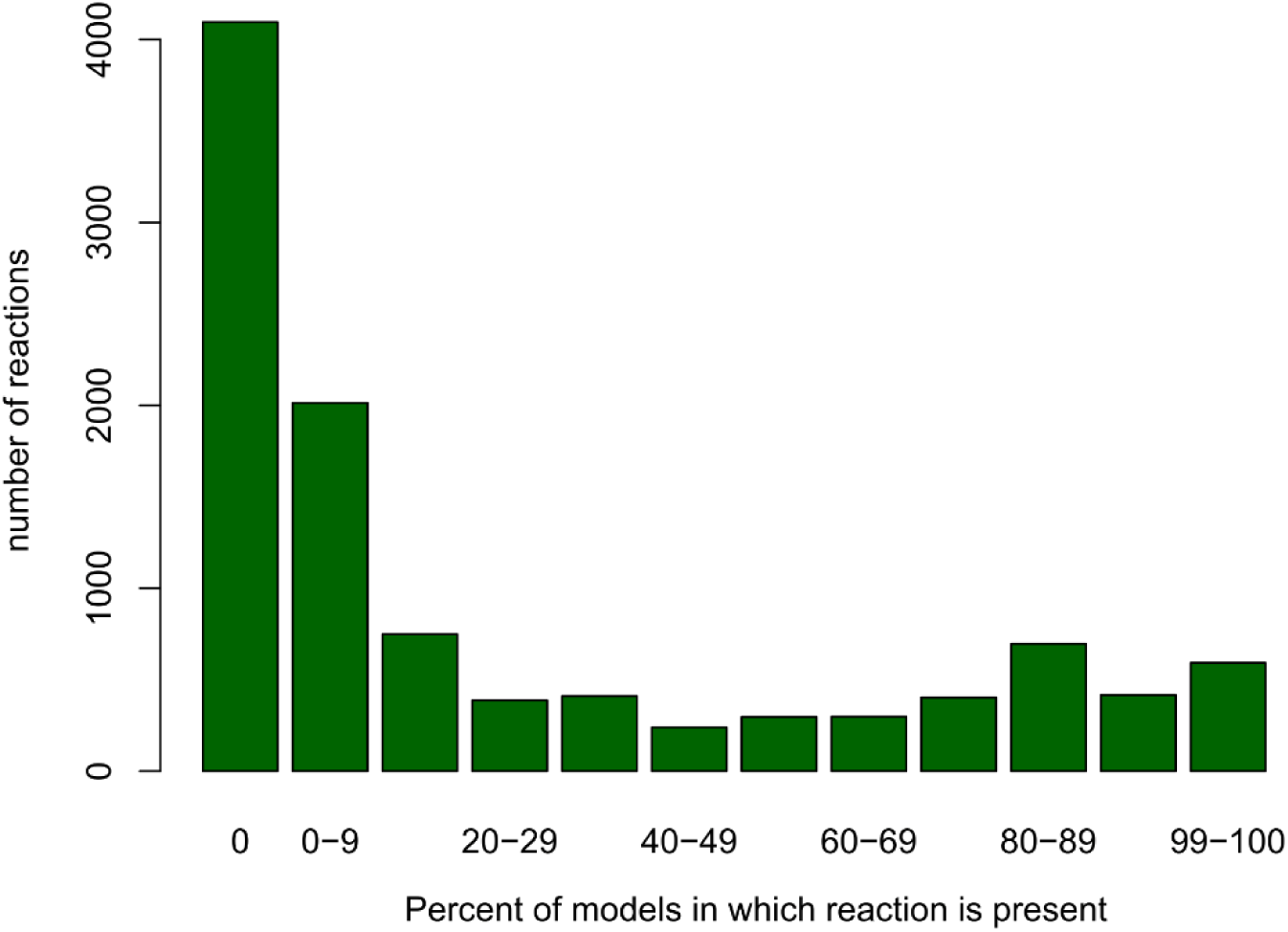
Distribution of the number of models in which each reaction is present across the 1,555 metabolic models.

**Supplementary Figure S3:**
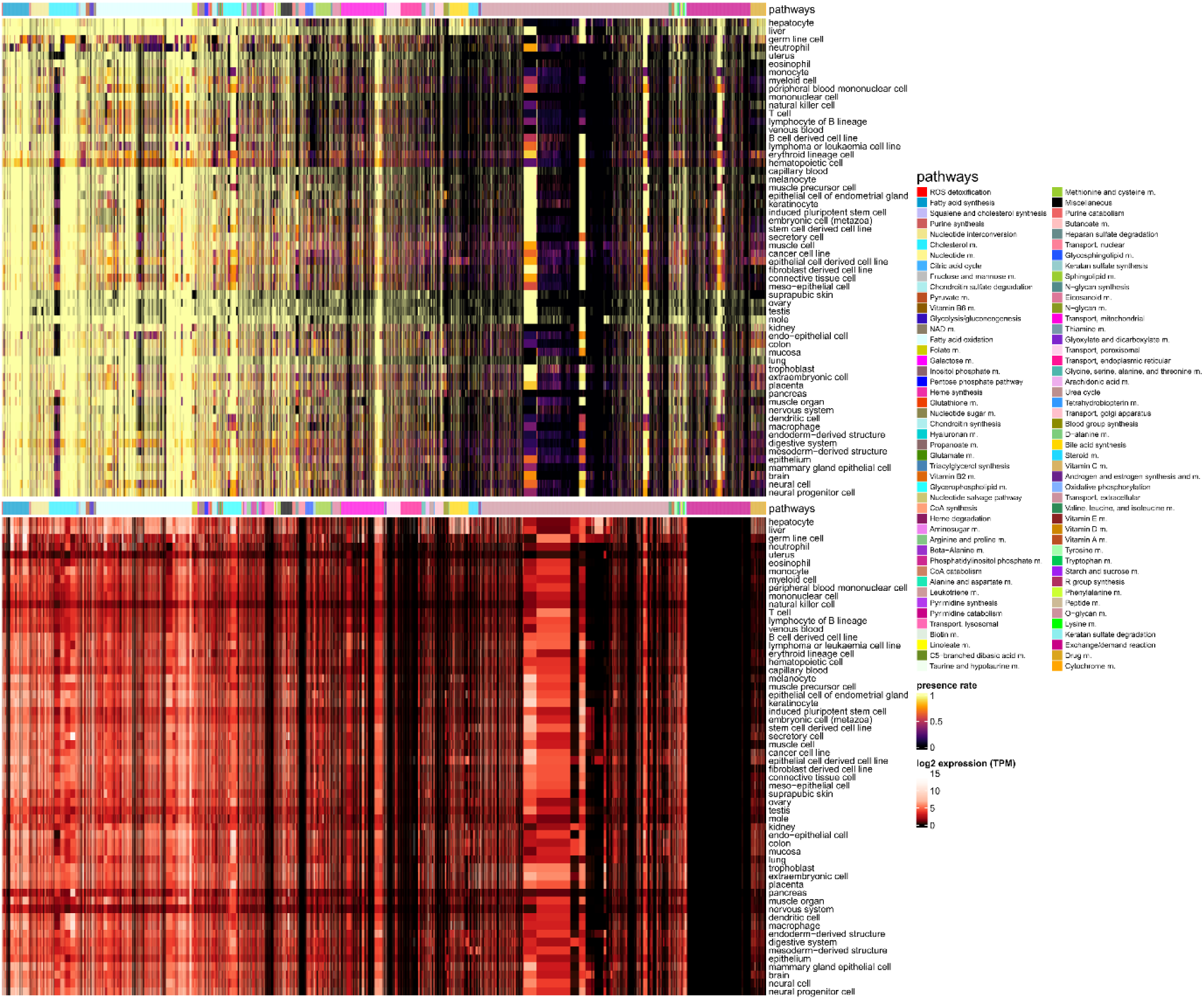
The presence rate of the reactions across the 1,555 models corresponds to the gene expression mapped to Recon3D via the Gene Protein Reaction rules.

**Supplementary Figure S4:**
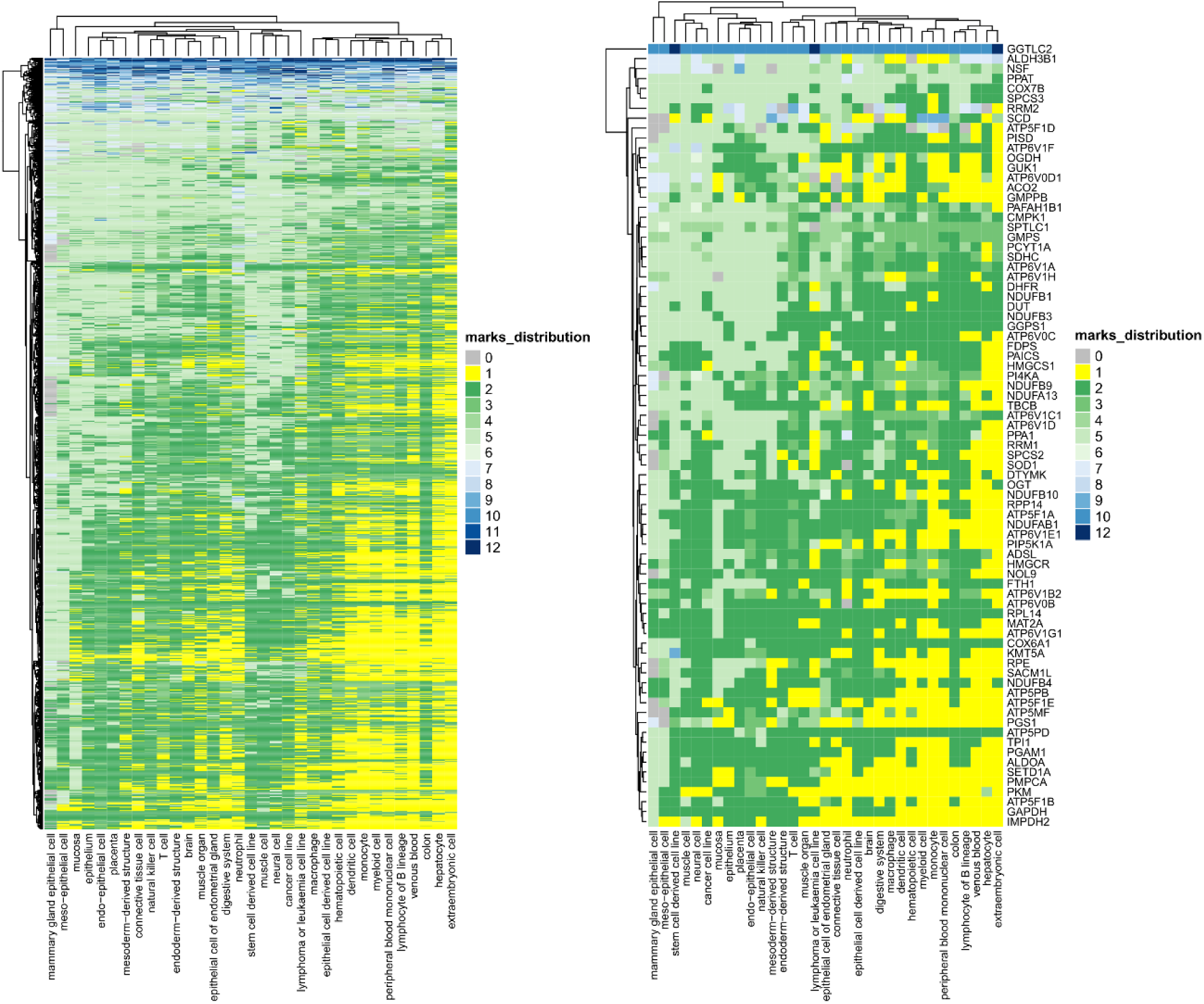
Metabolic gene ChromGene States across cell types. 60% of common (left) and metabolic essential genes (right) were in inactive ChromGene states (above seven, in blue) across 33 cell types. The remaining genes were tagged with active ChromGene state (1-6, yellow and green) for most cell types. However, several genes, such as gamma-glutamyl transferase (GGTLC2) were associated with repressive marks in almost all cell types.

**Supplementary Figure S5:**
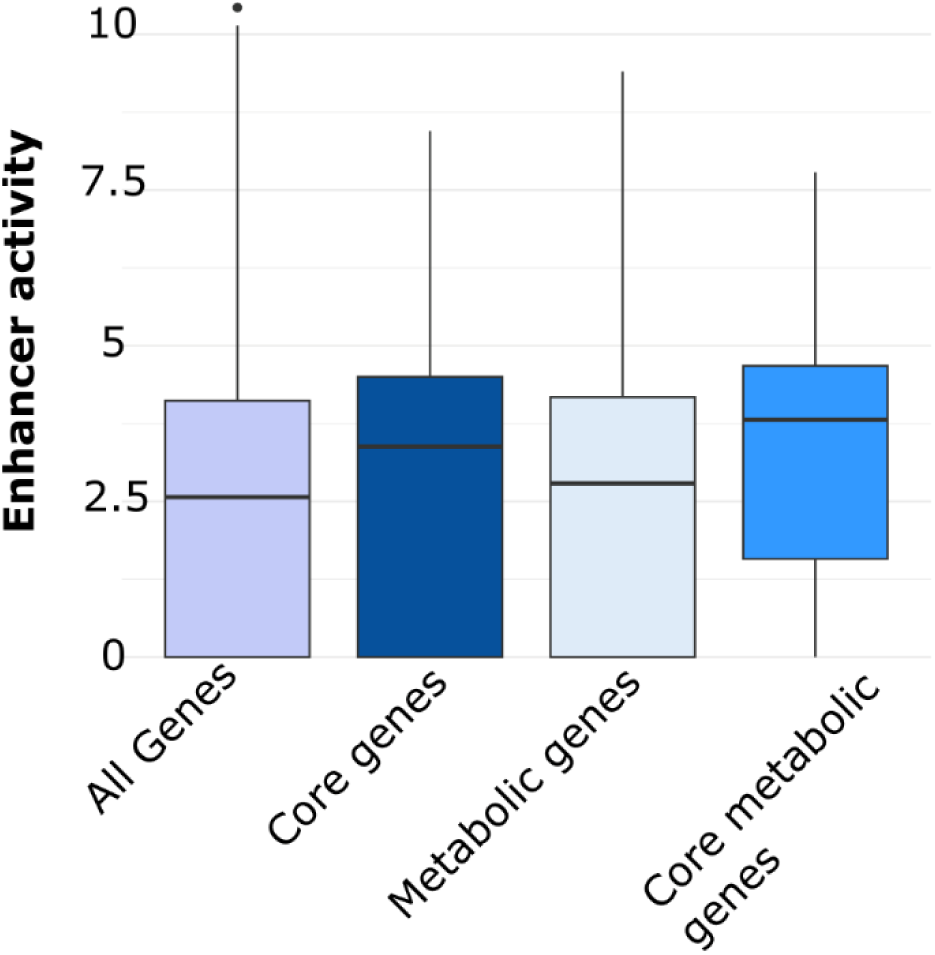
Unique genes and unique metabolic gene enhancer activities.

**Supplementary Figure S6:**
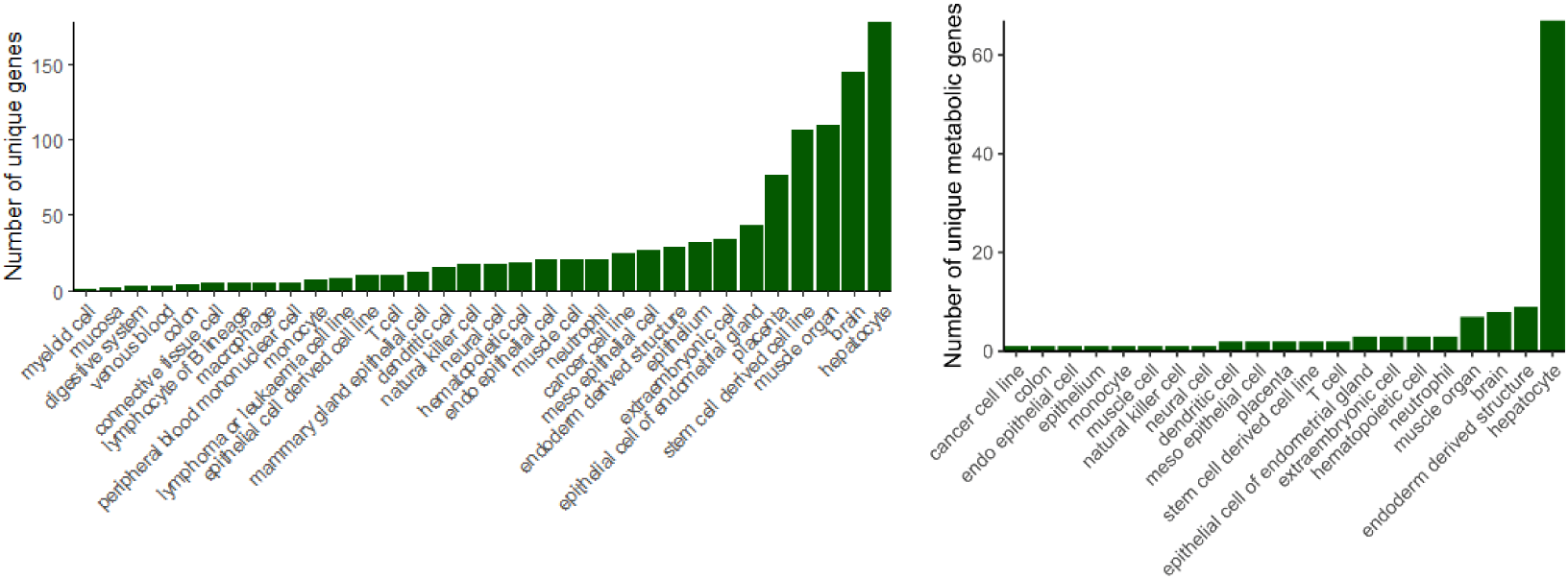
Repartition of the unique (left) and unique metabolic genes across tissue and cell types (right)

**Supplementary Figure S7:**
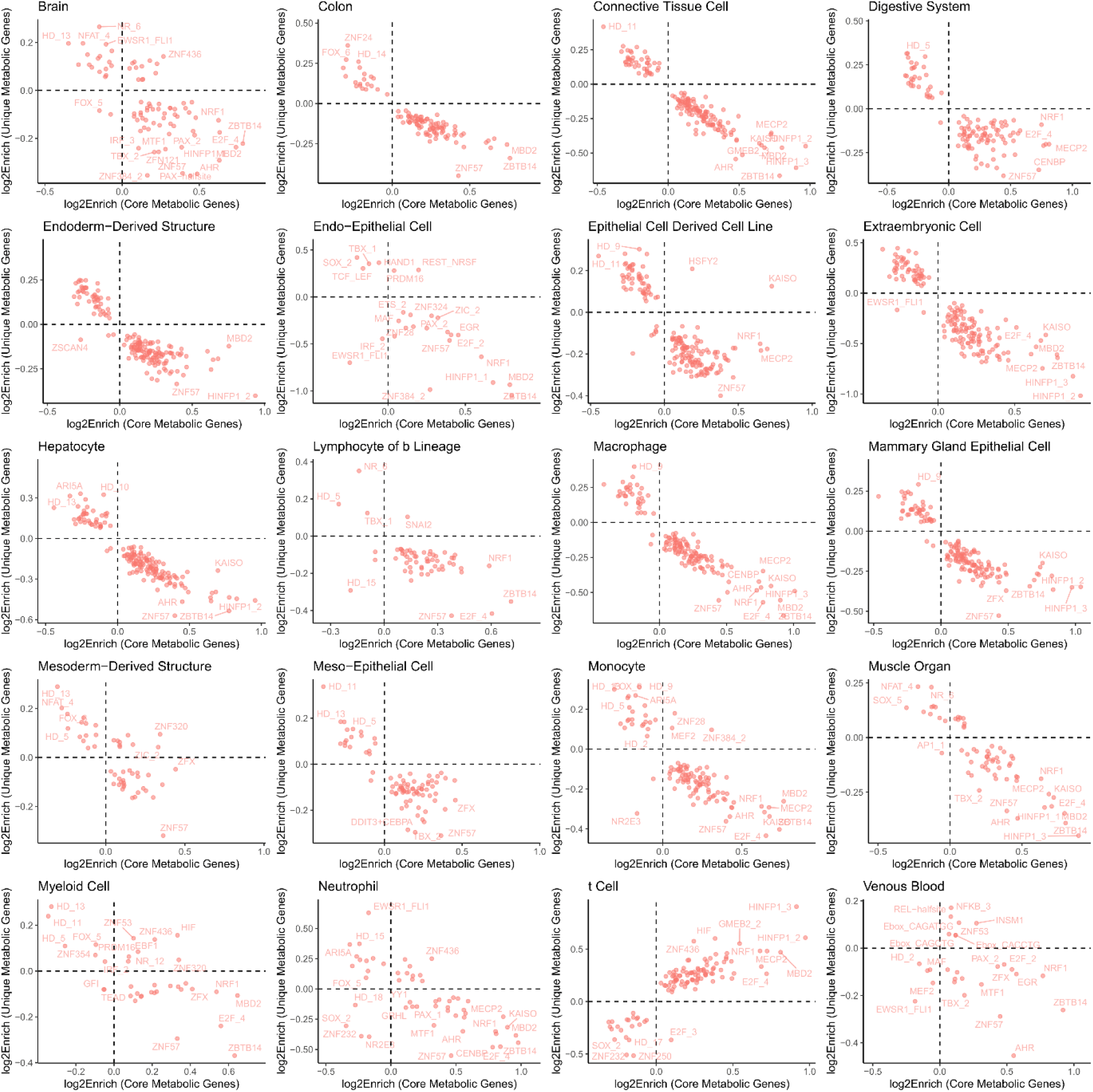
Correlation between TF enrichments in enhancers linked to unique metabolic genes vs core metabolic genes.

**Supplementary Figure S8:**
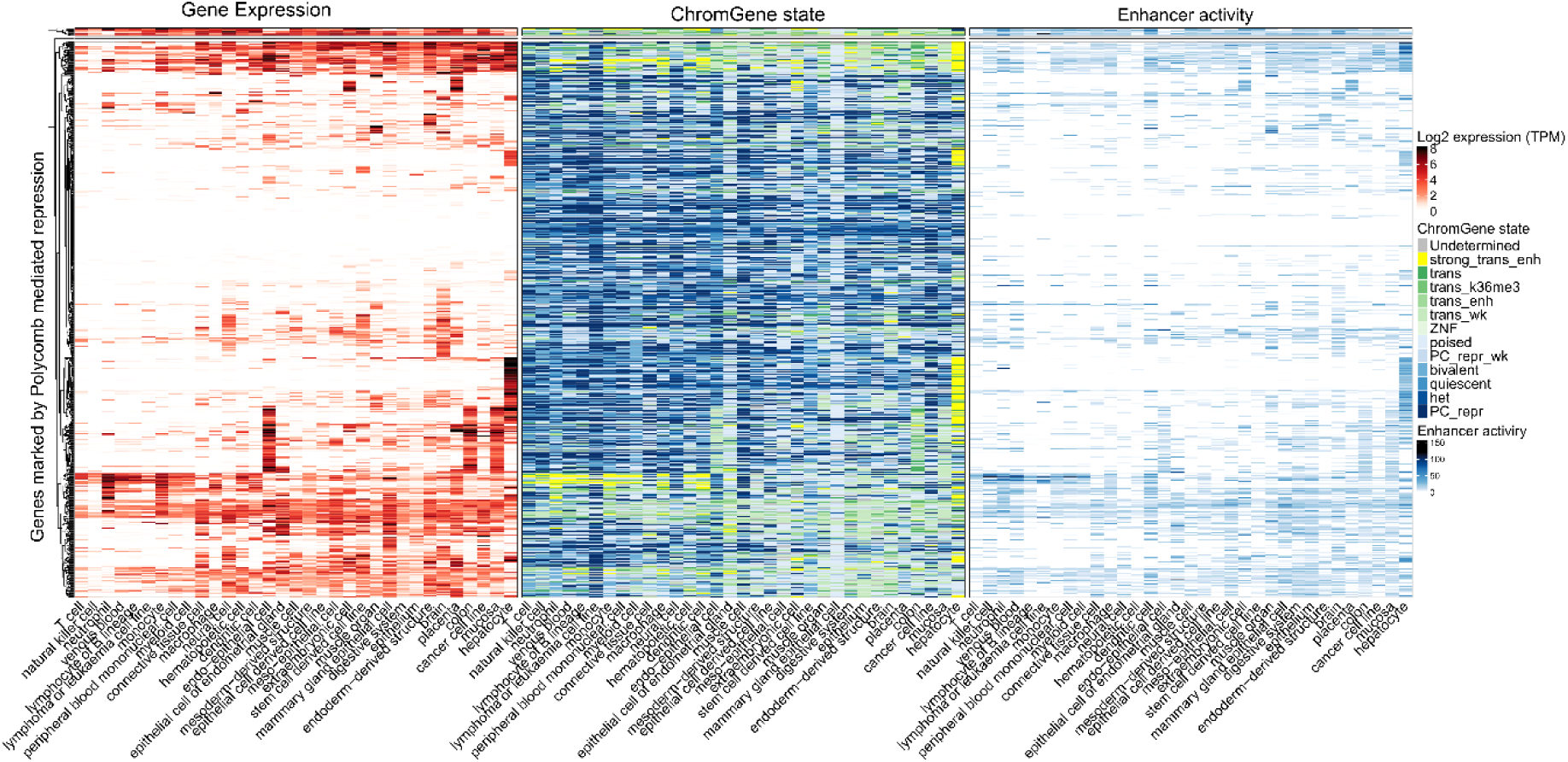
Expression, ChromGene states and sum enhancer and gene interactions for polycomb-associated genes.

**Supplementary Figure S9:**
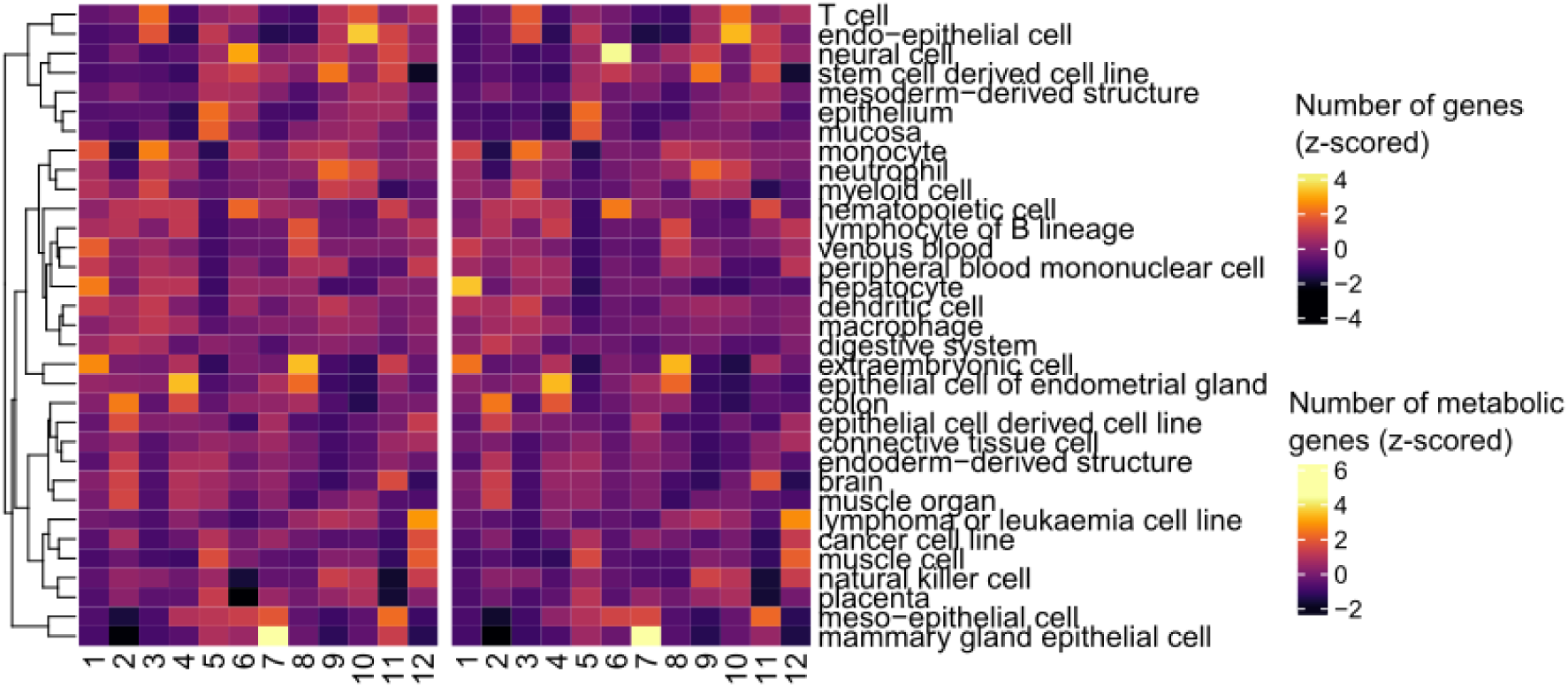
Number of genes associated with the twelve ChromGene states. Chrome 1-6 are linked to an active state, 7 stand for poised and 8-12 to repressive marks.

**Supplementary Figure S10:**
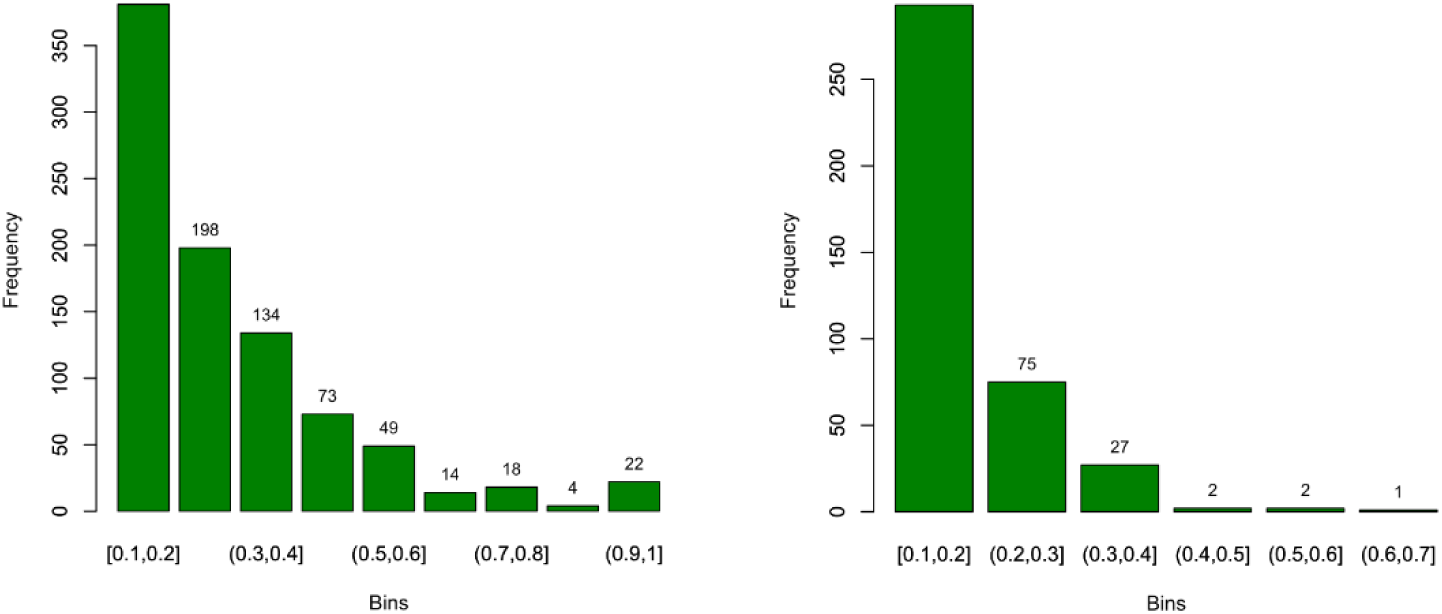
Number of pathway-cell type combinations having a presence rate between 0.1-0.7 for ChromGene 1 (left) and between 0.1-1 for ChromGene 12 (right).

**Supplementary Table S1:** The ChromGene state integrates active and repressive marks to obtain a gene-specific state instead of a locus state. 1-6 are active states, 7 corresponds to poised and 8-12 are repressive states.

| Nb | Name | Description |
| --- | --- | --- |
| 1 | strong trans enh | Strongly transcribed genes with enhancer marks H3K4me1 and H3K27ac, highest expressed |
| 2 | trans | Transcribed genes, highly expressed. Lower levels of H3K36me3 and H3K4me1 than strong trans enh |
| 3 | trans k36me3 | Transcribed genes, moderately expressed, marked with H3K36me3. Relatively low levels of enhancer marks |
| 4 | trans enh | Transcribed genes, moderately expressed, marked with enhancer marks H3K4me1 and to a lesser extent H3K27ac. Relatively low levels of H3K36me3. |
| 5 | trans wk | Transcribed genes, lowly expressed, marked with enhancer mark H3K27ac and to a lower extent, H3K4me3. Relatively low levels of H3K36me3 |
| 6 | ZNF | Zinc Finger-related genes, with a state marked by H3K36me3 and H3K9me3 |
| 7 | poised | Lowly-expressed genes with high levels of H3K4me1, but low levels of H3K36me3. |
| 8 | PC repr wk | Lowly expressed genes with high levels of H3K4me1, weakly marked with polycomb repressive mark H3K27me3. |
| 9 | bivalent | Lowly expressed genes, marked with high level of repressive mark H3K27me3, and moderate levels of H3K4me1 and H3K4me3 |
| 10 | quiescent | Lowly expressed genes, lack of significant histone marks |
| 11 | het | Generally unexpressed genes associated with heterochromatin. |
| 12 | PC repr | Generally, unexpressed genes, marked with polycomb repressive mark H3K27me3. |

**Supplementary Table S2:**
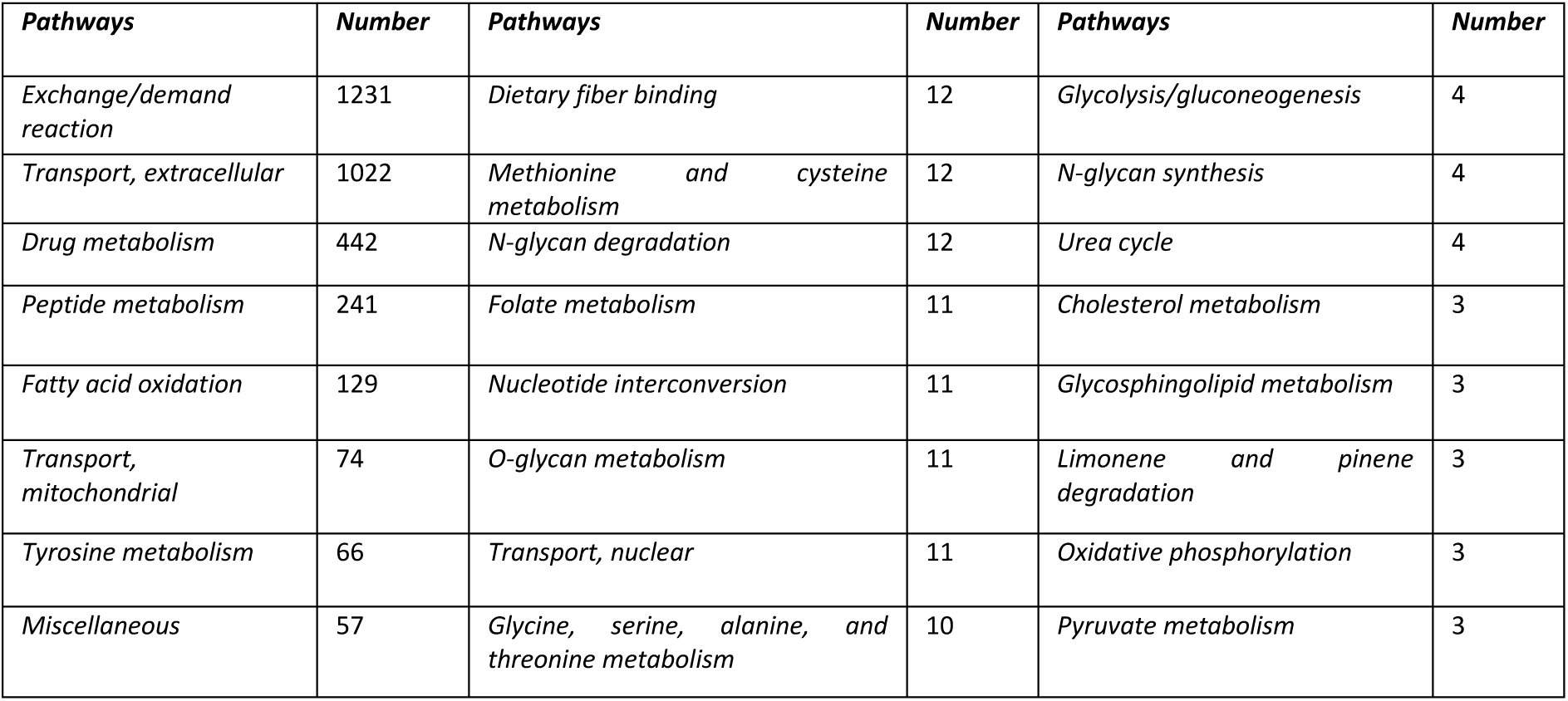
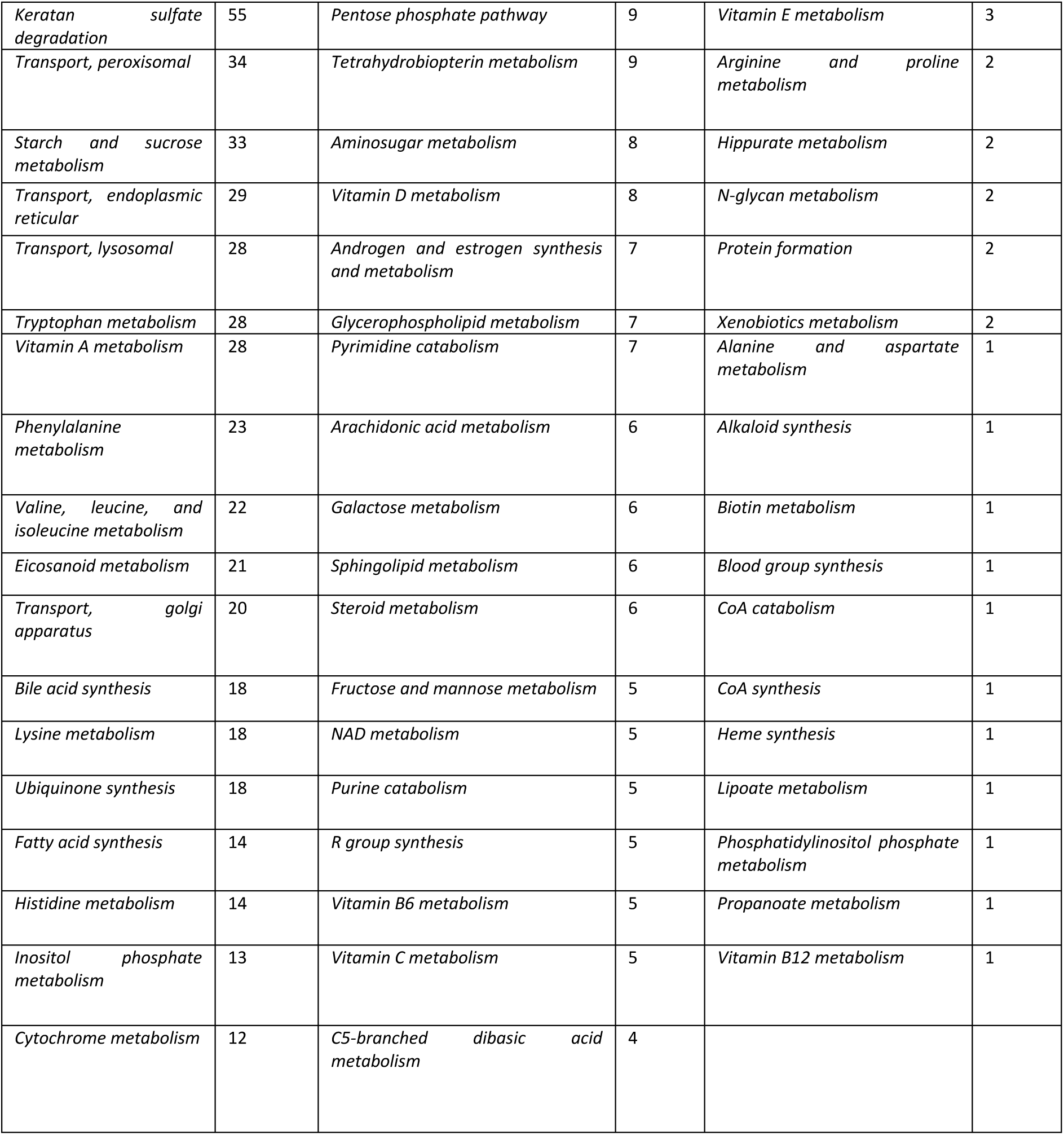
Number of reactions per pathway that were not included in any model.

**Supplementary Table S3:** Core metabolic genes are enriched for genes implicated in splicing and mRNA processing and energy production, nucleotide, lipid metabolism and ROS metabolism.

| GENE ONTOLOGY | OVERLAP | P-VALUE |
| --- | --- | --- |
| CORE GENES |  |  |
| MRNA SPLICING VIA SPLICEOSOME (GO:0000398) | 148/211 | 3.16E-60 |
| MRNA PROCESSING (GO:0006397) | 145/214 | 3.71E-56 |
| RNA SPLICING VIA TRANSESTERIFICATION REACTIONS WITH BULGED ADENOSINE AS NUCLEOPHILE (GO:0000377) | 130/180 | 1.77E-55 |
| TRANSLATION (GO:0006412) | 142/234 | 2.11E-46 |
| GENE EXPRESSION (GO:0010467) | 164/296 | 2.11E-46 |
| RIBOSOME BIOGENESIS (GO:0042254) | 106/155 | 1.66E-41 |
| RNA PROCESSING (GO:0006396) | 113/183 | 8.30E-38 |
| CYTOPLASMIC TRANSLATION (GO:0002181) | 74/93 | 3.65E-36 |
| RIBONUCLEOPROTEIN COMPLEX BIOGENESIS (GO:0022613) | 81/118 | 1.06E-31 |
| MACROMOLECULE BIOSYNTHETIC PROCESS (GO:0009059) | 104/183 | 2.35E-30 |
| RNA SPLICING (GO:0008380) | 71/98 | 4.03E-30 |
| PEPTIDE BIOSYNTHETIC PROCESS (GO:0043043) | 93/158 | 9.88E-29 |
| PROTEIN-RNA COMPLEX ASSEMBLY (GO:0022618) | 89/150 | 7.60E-28 |
| RIBOSOMAL SMALL SUBUNIT BIOGENESIS (GO:0042274) | 61/84 | 6.70E-26 |
| RRNA PROCESSING (GO:0006364) | 68/101 | 7.91E-26 |
| PROTEASOME-MEDIATED UBIQUITIN-DEPENDENT PROTEIN CATABOLIC PRO. | 139/319 | 4.21E-25 |
| NCRNA PROCESSING (GO:0034470) | 63/100 | 1.63E-21 |
| RRNA METABOLIC PROCESS (GO:0016072) | 59/91 | 4.69E-21 |
| METABOLIC CORE GENES |  |  |
| METABOLISM R-HSA-1430728 | 210/2049 | 6.61E-133 |
| CITRIC ACID (TCA) CYCLE AND RESPIRATORY ELECTRON TRANSPORT | 64/163 | 7.18E-72 |
| RESPIRATORY ELECTRON TRANSPORT, ATP SYNTHESIS BY CHEMIOSMOTIC COUPLING, HEAT PRODUCTION BY UNCOUPLING PROTEINS R-HSA-163200 | 53/112 | 1.35E-64 |
| RESPIRATORY ELECTRON TRANSPORT R-HSA-611105 | 44/90 | 3.55E-54 |
| COMPLEX I BIOGENESIS R-HSA-6799198 | 27/51 | 4.69E-34 |
| METABOLISM OF NUCLEOTIDES R-HSA-15869 | 24/95 | 6.30E-21 |
| METABOLISM OF LIPIDS R-HSA-556833 | 51/732 | 4.91E-18 |
| CHOLESTEROL BIOSYNTHESIS R-HSA-191273 | 11/26 | 3.32E-12 |
| INTERCONVERSION OF NUCLEOTIDE DI- AND TRIPHOSPHATES R-HSA-499943 | 11/28 | 7.98E-12 |
| METABOLISM OF CARBOHYDRATES R-HSA-71387 | 26/285 | 1.52E-11 |
| FORMATION OF ATP BY CHEMIOSMOTIC COUPLING R-HSA-163210 | 9/16 | 1.81E-11 |
| PURINE RIBONUCLEOSIDE MONOPHOSPHATE BIOSYNTHESIS R-HSA-73817 | 7/10 | 8.26E-10 |
| PYRUVATE METABOLISM AND CITRIC ACID (TCA) CYCLE R-HSA-71406 | 12/54 | 9.25E-10 |
| NUCLEOTIDE BIOSYNTHESIS R-HSA-8956320 | 7/13 | 9.74E-09 |
| CRISTAE FORMATION R-HSA-8949613 | 9/29 | 9.78E-09 |
| ACTIVATION OF GENE EXPRESSION BY SREBF (SREBP) R-HSA-2426168 | 10/42 | 1.66E-08 |
| METABOLISM OF VITAMINS AND COFACTORS R-HSA-196854 | 18/186 | 1.66E-08 |
| FATTY ACID METABOLISM R-HSA-8978868 | 17/173 | 3.74E-08 |
| PKMTS METHYLATE HISTONE LYSINES R-HSA-3214841 | 10/47 | 4.70E-08 |
| METABOLISM OF WATER-SOLUBLE VITAMINS AND COFACTORS R-HSA-196849 | 14/122 | 1.30E-07 |
| REGULATION OF CHOLESTEROL BIOSYNTHESIS BY SREBP (SREBF) R-HSA-1655829 | 10/55 | 2.16E-07 |
| GLUCONEOGENESIS R-HSA-70263 | 8/32 | 4.47E-07 |
| BIOSYNTHESIS OF N-GLYCAN PRECURSOR (DOLICHOL LLO) AND TRANSFER TO PROTEIN R-HSA-446193 | 11/77 | 4.99E-07 |
| HEME BIOSYNTHESIS R-HSA-189451 | 6/14 | 6.47E-07 |
| SYNTHESIS OF SUBSTRATES IN N-GLYCAN BIOSYTHESIS R-HSA-446219 | 10/63 | 7.11E-07 |
| TP53 REGULATES METABOLIC GENES R-HSA-5628897 | 11/81 | 7.59E-07 |
| BIOLOGICAL OXIDATIONS R-HSA-211859 | 17/218 | 8.13E-07 |
| PHASE II - CONJUGATION OF COMPOUNDS R-HSA-156580 | 12/107 | 1.51E-06 |
| MITOCHONDRIAL BIOGENESIS R-HSA-1592230 | 11/89 | 1.84E-06 |
| METABOLISM OF PORPHYRINS R-HSA-189445 | 7/28 | 2.57E-06 |
| CYTOPROTECTION BY HMOX1 R-HSA-9707564 | 9/57 | 2.92E-06 |
| IRON UPTAKE AND TRANSPORT R-HSA-917937 | 9/59 | 3.85E-06 |
| PYRUVATE METABOLISM R-HSA-70268 | 7/31 | 5.00E-06 |
| METABOLISM OF STEROIDS R-HSA-8957322 | 13/153 | 9.50E-06 |
| MITOCHONDRIAL FATTY ACID BETA-OXIDATION R-HSA-77289 | 7/36 | 1.40E-05 |
| INSULIN RECEPTOR RECYCLING R-HSA-77387 | 6/26 | 2.83E-05 |
| SYNTHESIS OF UDP-N-ACETYL-GLUCOSAMINE R-HSA-446210 | 4/8 | 4.54E-05 |
| VITAMIN B5 (PANTOTHENATE) METABOLISM R-HSA-199220 | 5/17 | 5.33E-05 |
| CELLULAR RESPONSE TO CHEMICAL STRESS R-HSA-9711123 | 13/182 | 5.80E-05 |
| GLYCOSPHINGOLIPID METABOLISM R-HSA-1660662 | 7/45 | 5.92E-05 |
| TRANSFERRIN ENDOCYTOSIS AND RECYCLING R-HSA-917977 | 6/31 | 7.45E-05 |
| ASPARAGINE N-LINKED GLYCOSYLATION R-HSA-446203 | 16/282 | 8.72E-05 |
| SPHINGOLIPID METABOLISM R-HSA-428157 | 9/89 | 9.47E-05 |
| GLUCOSE METABOLISM R-HSA-70326 | 9/89 | 9.47E-05 |

**Supplementary Table S4:**
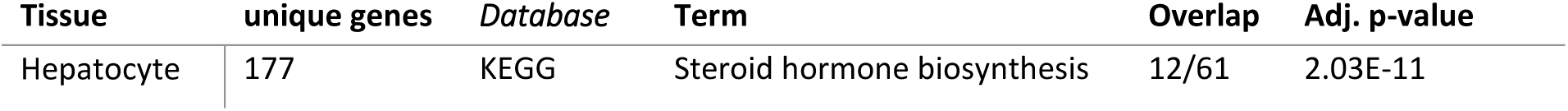
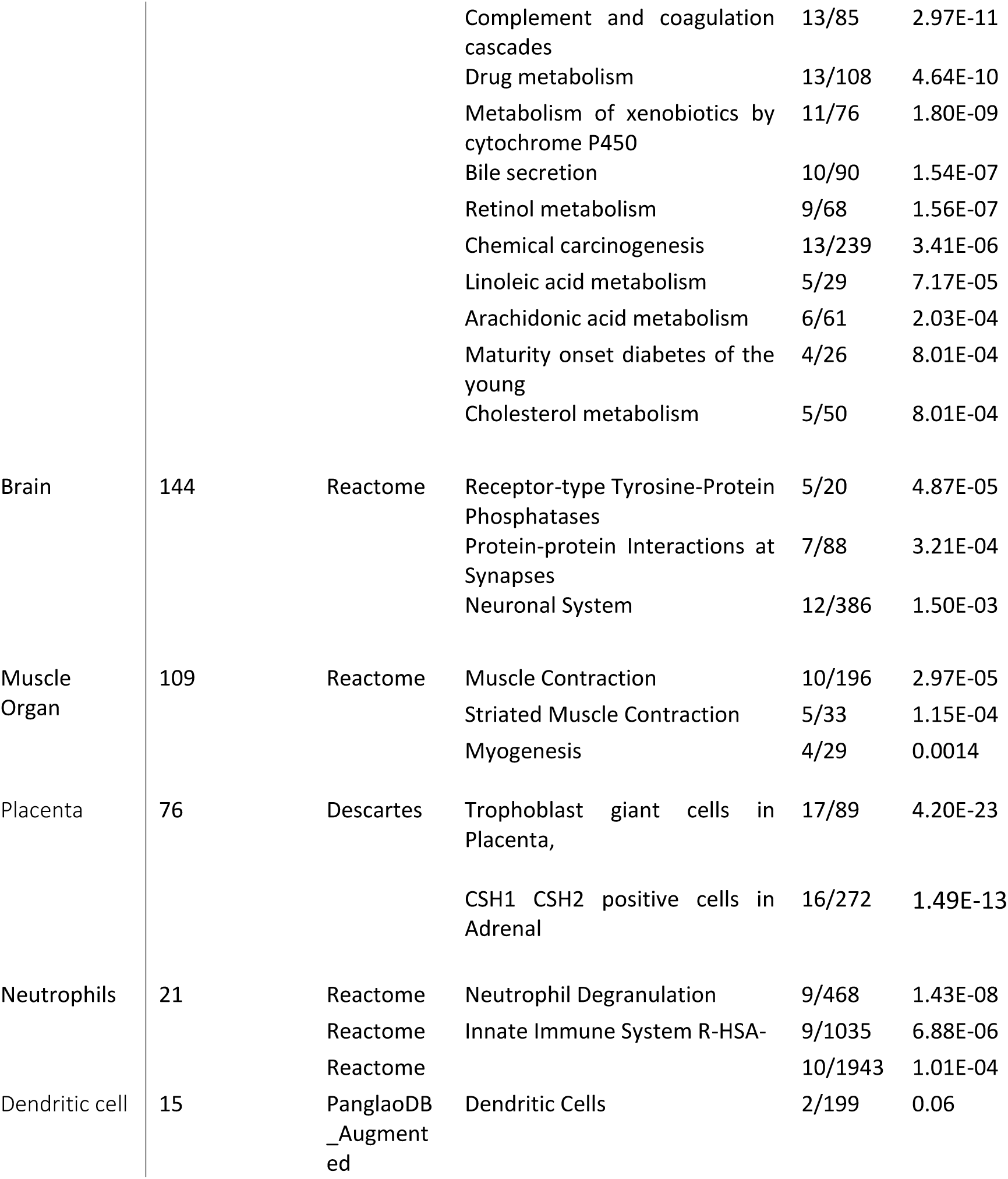
Unique genes for hepatocyte, brain, and muscle organs are enriched for organ-matched gene ontology terms. Unique genes were defined by having an active consensus ChromGene state (states below 7) in one single tissue or cell type. No significant enrichments were found for colon and connective tissue cell line due to a limited number of genes (4, 5 genes). Similarly, no enrichments were found for cancer lines due to the substantial tissue heterogeneity of this category.

**Supplementary Table S5:**
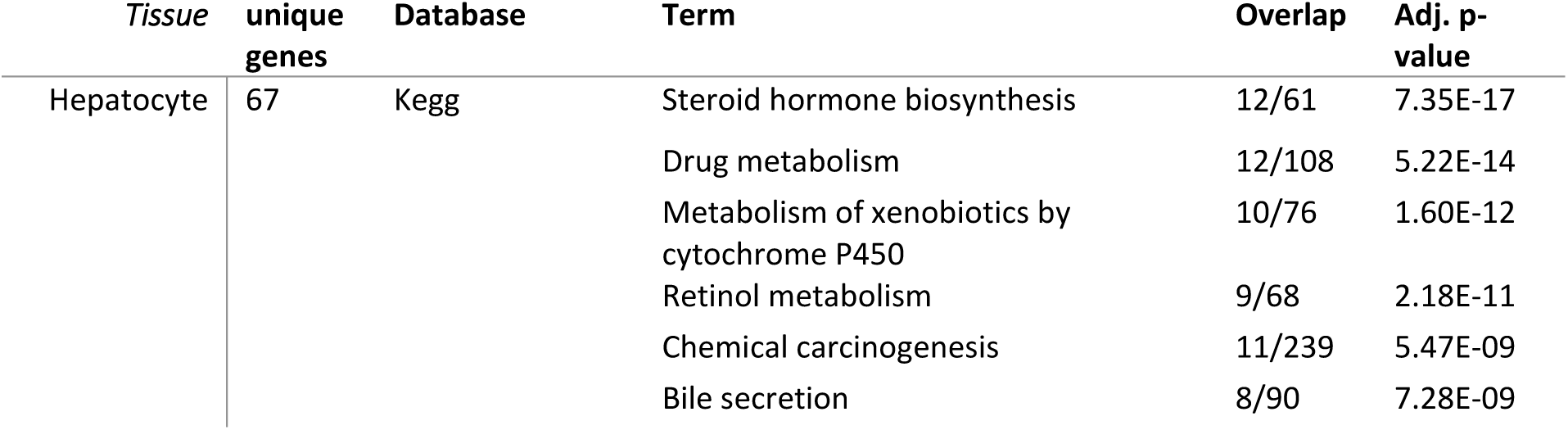
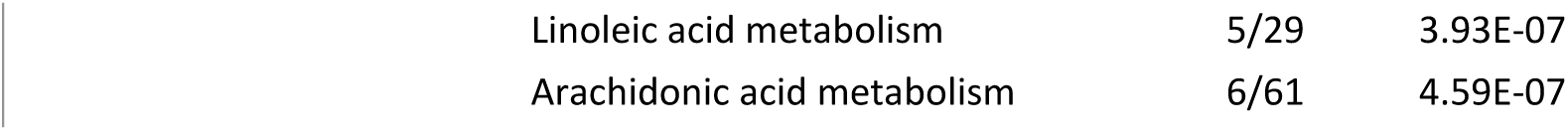
Unique metabolic genes for hepatocyte, brain, and muscle are enriched for organ-matched gene ontology terms. Unique genes specific to one tissue were defined by having a consensus ChromGene below 7 in one single tissue or cell type.

**Supplementary Table S8:**
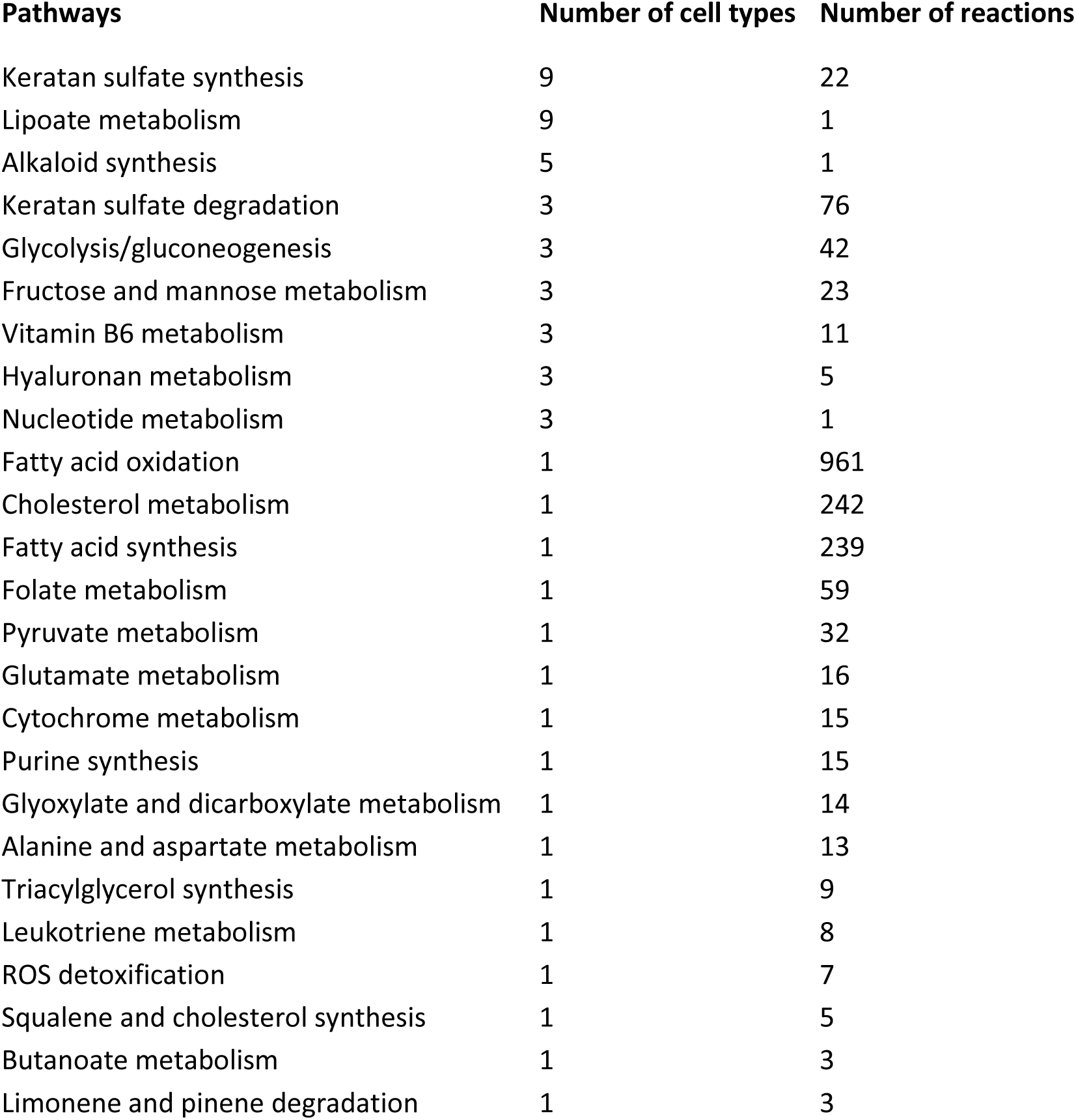
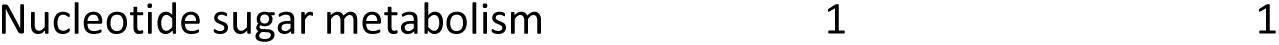
Number of cell types with a presence rate above 0.6 for strongly transcribed genes with enhancer marks H3K4me1 and H3K27ac (ChromGene State 1)

**Supplementary Table S9:** Number of cell types with a presence rate above 0.3 for Polycomb-mediated.

| **Pathways** | **Number of cell types** | **Number of reactions** |
| --- | --- | --- |
| Cytochrome metabolism | 12 | 15 |
| Vitamin D metabolism | 4 | 13 |
| Alanine and aspartate metabolism | 3 | 13 |
| Steroid metabolism | 3 | 90 |
| Bile acid synthesis | 2 | 185 |
| D-alanine metabolism | 2 | 3 |
| Androgen and estrogen synthesis and metabolism | 1 | 26 |
| Blood group synthesis | 1 | 47 |
| Butanoate metabolism | 1 | 3 |
| Glycine, serine, alanine, and threonine metabolism | 1 | 47 |
| O-glycan metabolism | 1 | 18 |
| Triacylglycerol synthesis | 1 | 9 |

